# Cbln1 regulates axon growth and guidance in multiple neural regions

**DOI:** 10.1101/2021.11.16.468844

**Authors:** Peng Han, Yuanchu She, Zhuoxuan Yang, Mengru Zhuang, Qingjun Wang, Xiaopeng Luo, Chaoqun Yin, Junda Zhu, Samie R. Jaffrey, Sheng-Jian Ji

## Abstract

The accurate construction of neural circuits requires the precise control of axon growth and guidance, which is regulated by multiple growth and guidance cues during early nervous system development. It is generally thought that the growth and guidance cues that control the major steps of axon guidance have been defined. Here, we describe cerebellin-1 (Cbln1) as a novel cue that controls diverse aspects of axon growth and guidance throughout the central nervous system (CNS). Cbln1 has previously been shown to function in late neural development to influence synapse organization. Here we find that Cbln1 has an essential role in early neural development. Cbln1 is expressed on the axons and growth cones of developing commissural neurons and functions in an autocrine manner to promote axon growth. Cbln1 is also expressed in intermediate target tissues and functions as an attractive guidance cue. We find that these functions of Cbln1 are mediated by neurexin-2 (Nrxn2), which functions as the Cbln1 receptor for axon growth and guidance. In addition to the developing spinal cord, we further show that Cbln1 functions in diverse parts of the CNS with major roles in cerebellar parallel fiber growth and retinal ganglion cell axon guidance. Despite the prevailing role of Cbln1 as a synaptic organizer, our study discovers a new and unexpected function for Cbln1 as a general axon growth and guidance cue throughout the nervous system.

**Impact statement:** Despite the prevailing role of Cbln1 as a synaptic organizer, our study discovers a new and unexpected function for Cbln1 as a general axon growth and guidance cue throughout the nervous system.

## Introduction

The precise control of axon pathfinding is critical for the correct neural wiring during nervous system development. The stimulation of axon growth and regulation of axon guidance have been shown to require adhesion molecules, diffusible signals and morphogens such as Netrins ***(Moreno-Bravo et al. 2019; Wu et al. 2019)***, Slits ***(Brose et al. 1999; Kidd et al. 1999; Zou et al. 2000)***, Ephrins ***(Paixao et al. 2013)***, Semaphorins ***(Zou et al. 2000; Nawabi et al. 2010)***, Draxin ***(Islam et al. 2009)***, Shh ***(Okada et al. 2006)***, Wnts ***(Lyuksyutova et al. 2003)***, and BMPs ***(Augsburger et al. 1999; Butler and Dodd 2003)***. These axon guidance molecules bind to their receptors in the axon growth cones to activate various signaling pathways that eventually change the cytoskeleton ***(McCormick and Gupton 2020)***. The lack of newly identified cues in the past decade has suggested that the major classes of growth and guidance cues have now been identified.

The commissures in the rodent spinal cord are one of the most prominent model systems to study axon growth and guidance. In a search for the differentially expressed genes in the dorsal spinal cord of mouse embryos, we identified a gene encoding the secreted protein cerebellin-1 (Cbln1). Cbln1 is released from cerebellar parallel fibers and has previously been characterized as a synaptic organizer by forming the synapse-spanning tripartite complex Nrxn-Cbln1-GluD2 (Nrxn, neurexin; GluD2, the ionotropic glutamate receptor family member delta-2) ***(Yuzaki 2018; Suzuki et al. 2020)***. However, whether Cbln1 is expressed and plays roles in earlier nervous system development is unknown.

Here we found that Cbln1 is expressed both in the dorsal commissural neurons (DCN) and in the floor plate (FP) of the embryonic mouse spinal cord. We generated DCN- and FP-specific *Cbln1* conditional knockout (cKO) mice which demonstrated that the cell-autonomous and non-cell-autonomous Cbln1 from DCNs and FP regulate commissural axon growth and guidance, respectively. The dual roles of Cbln1 are mediated by its receptor, neurexin-2. Interestingly, the functions and mechanisms of Cbln1 in regulating axon growth and guidance were replicated in the developing cerebellar granule axon growth and the embryonic retinal ganglion cell axon guidance, respectively.

Together, our findings reveal a general role for Cbln1 in regulating axon growth and guidance during early nervous system development prior to synapse formation.

## Results

### Cbln1 is expressed in both dorsal commissural neurons and floor plate in the developing mouse spinal cord

To identify the differentially expressed genes in the mouse embryonic dorsal spinal cord, we genetically labeled embryonic dorsal spinal neurons with eGFP by crossing *Wnt1-cre **(Danielian et al. 1998; Charron et al. 2003)*** with *Rosa26mTmG **(Muzumdar et al. 2007)*** mice (*Figure 1—figure supplement 1A*). Mouse embryonic E10.5, E11.5 and E12.5 spinal cords were dissected, and dorsal spinal neurons were collected (*Figure 1—figure supplement 1B*). Then GFP^+^ dorsal spinal neurons were purified by fluorescence-activated cell sorting (FACS) and the differentially expressed genes (DEGs) during the developmental stages were identified by the expression profiling analysis using microarray analysis (*Figure 1—figure supplement 1C* and *Supplementary file 1*). We carried out *in situ* hybridization to further explore the expression patterns of the candidate DEGs in the developing spinal cord. Among the candidates, *Cbln1* was notable due to its expression pattern. *Cbln1* has strong signals in floor plate (FP) and weak signals in dorsal commissural neurons (DCNs) at E10.5 (*Figure 1A*). At E11.5 and E12.5, the expression of *Cbln1* increases in DCNs (notice DCNs migrate ventrally and medially at E12.5), maintains a high level in FP, and also appears in subpopulations of motor neurons (*Figure 1A*).

**Figure 1.**
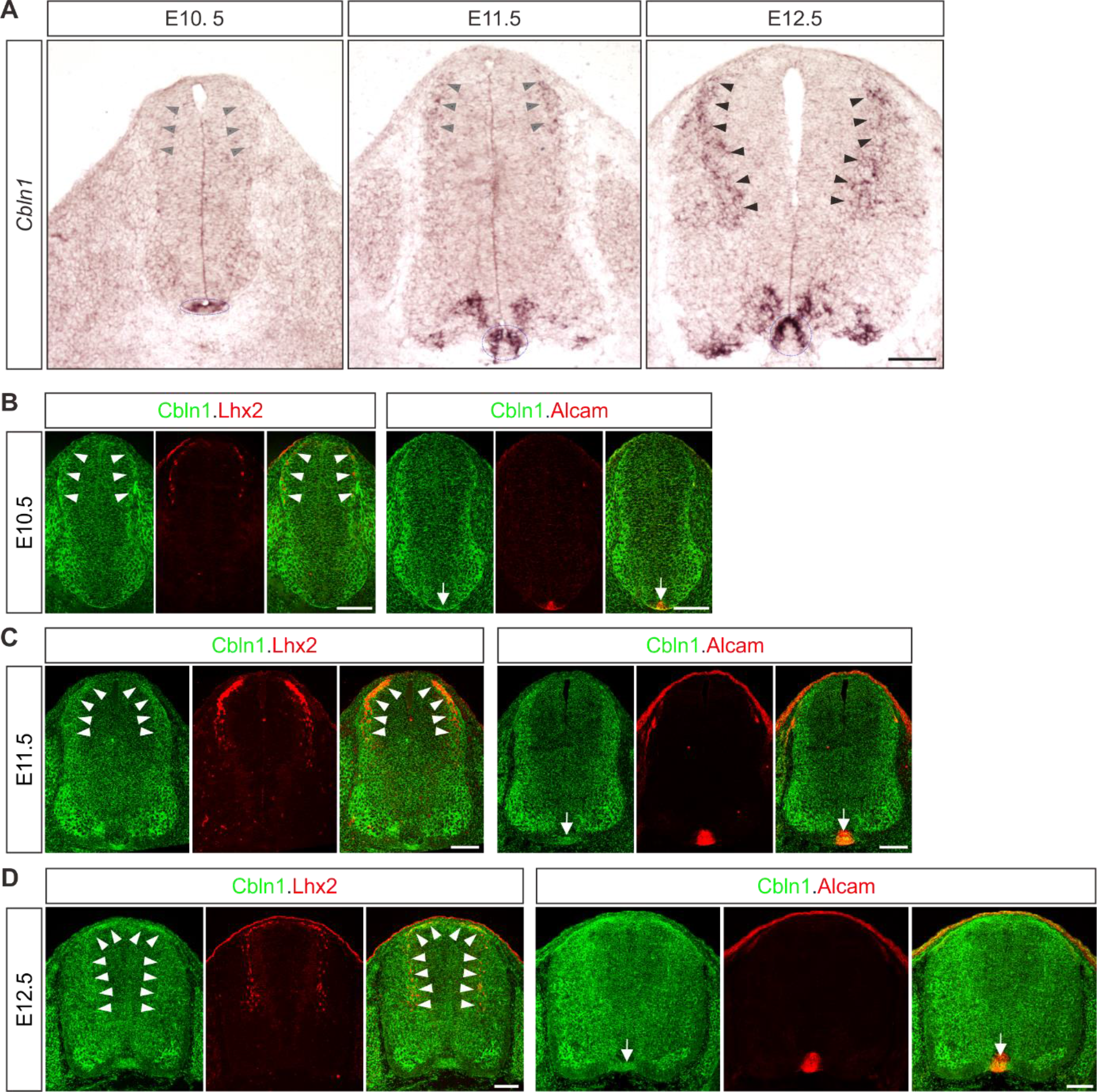
Expression patterns of *Cbln1* and Cbln1 in the developing mouse spinal cord. **(A)** *In situ* hybridization was carried out using a DIG-labelled RNA probe against *Cbln1* in spinal cord cross-sections during E10.5∼E12.5. Arrowheads and circled areas indicate the expression of *Cbln1* in the dorsal commissural neurons (DCN) and the floor plate, respectively. Scale bar, 100 μm. **(B-D)** Co-immunostaining of Cbln1 with Lhx2 or Alcam in spinal cord cross-sections during E10.5∼E12.5 showed expression of Cbln1 in the dorsal commissural neurons and floor plate. White arrowheads and arrows point to the expression of Cbln1 in the dorsal commissural neurons and floor plate, respectively. Scale bars, 100 μm.

To further explore the expression patterns of Cbln1 protein and confirm its expression sites, we carried out immunofluorescence (IF) using a Cbln1 antibody ***(Muguruma et al. 2010)***. Co-immunostaining of Cbln1 with Lhx2, a DCN marker ***(Wilson et al. 2008)***, confirmed the expression of Cbln1 in DCNs of developing spinal cords from E10.5 to E12.5 (*Figure 1B-D*). Expression of Cbln1 in FP was also confirmed by co-immunostaining with the FP marker Alcam (*Figure 1B-D*). To validate the specificity of Cbln1 expression in FP, we used a spinal floor plate-deficient model, *Gli2* knockout (KO) mouse ***(Bai and Joyner 2001)***. As shown in *Figure 1—figure supplement 1D*, IF signal of Cbln1 in Alcam-marked FP was gone in *Gli2* KO. These results revealed an interesting expression pattern for Cbln1 that is expressed both in the dorsal commissural neurons (DCNs) and in the intermediate target for DCNs, the floor plate.

### Cell-autonomous Cbln1 in the dorsal commissural neurons stimulates commissural axon growth in an autocrine manner

Next, we wanted to explore the roles of Cbln1 expressed in DCNs and floor plate, separately. In order to specifically ablate *Cbln1* from these tissues, we generated conditional knockouts (cKO) of *Cbln1* using tissue-specific *Cre* lines (*Figure 2—figure supplement 1A*). We used *Wnt1-Cre* line to specifically ablate *Cbln1* from spinal DCNs, without affecting *Cbln1* expression in other parts of spinal cord (*Figure 2A*). *Cbln*1 cKO in spinal DCNs does not disturb neurogenesis of these neurons, as indicated by normal numbers, distribution and patterning of Lhx2^+^ and Lhx9^+^ interneurons in the developing spinal cord (*Figure 2—figure supplement 1B-D*). We continued to check commissural axon (CA) growth in DCN-specific *Cbln1* cKO. We prepared open-books of developing spinal cords and used Robo3 immunostaining to label commissural axons. Robo3 selectively marks commissural axons as they navigate to and across the floor plate ***(Sabatier et al. 2004)***. As shown in *Figure 2B-D*, both lengths and numbers of commissural axons were decreased in *Cbln1* cKO embryos compared with their littermate controls. These data suggest that Cbln1 in the dorsal commissural neurons is required for their own commissural axon growth.

**Figure 2.**
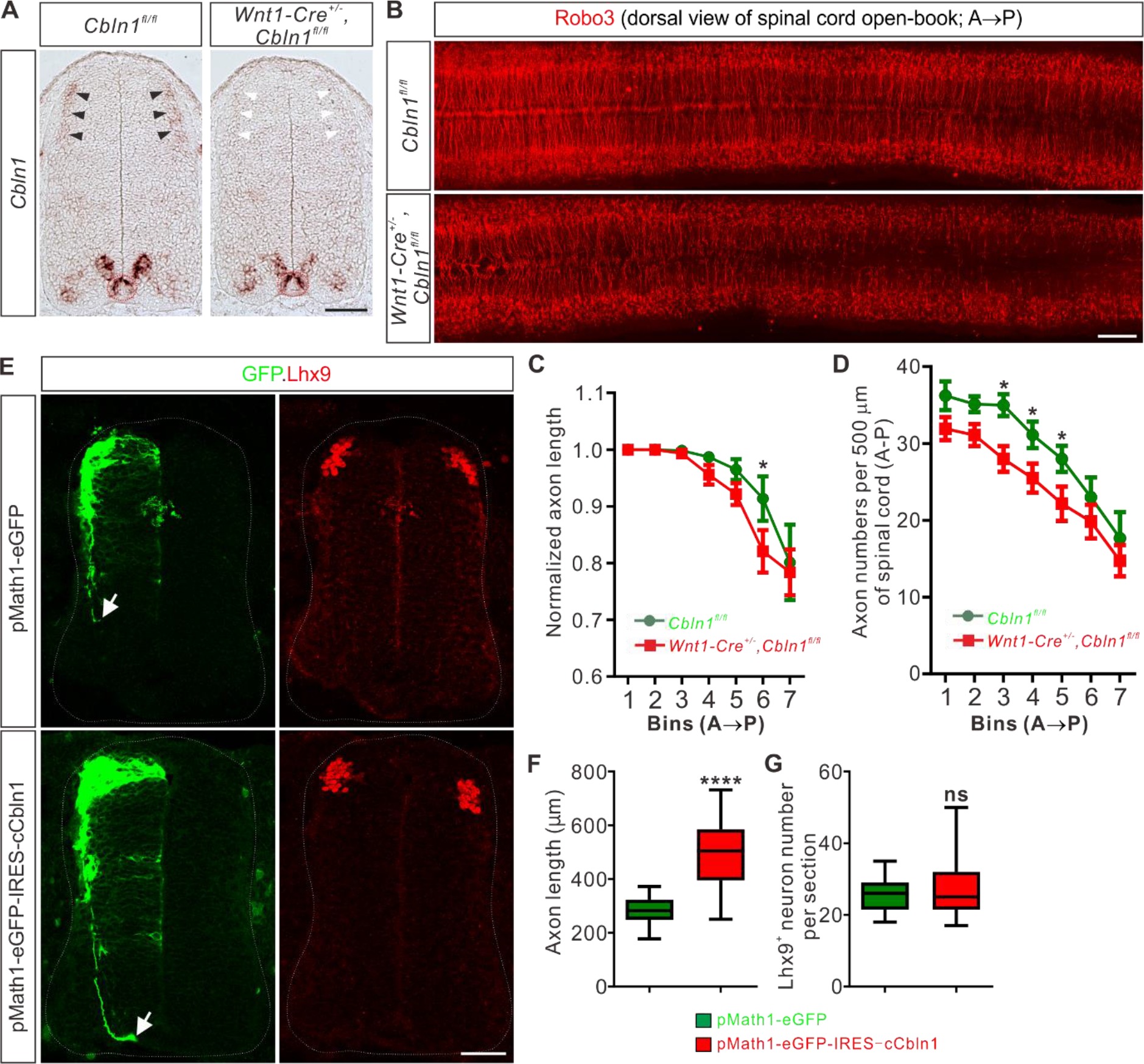
Cell-autonomous Cbln1 is both required and sufficient to stimulate commissural axon growth *in vivo*. **(A)** Specific depletion of *Cbln1* in the dorsal spinal cord of *Wnt1-Cre^+/-^,Cbln1^fl/fl^* cKO mouse embryos. *In situ* hybridization of E11.5 spinal cord sections using RNA probes against *Cbln1* confirmed specific ablation of *Cbln1* from the dorsal commissural neurons (DCNs). Black arrowheads indicate *Cbln1* expression in control DCNs and white arrowheads highlight the missing *Cbln1* expression in cKO DCNs. Expression of *Cbln1* in floor plate (in red dotted circles) and other parts are not affected in DCN-specific *Cbln1* cKO spinal cords. Scale bar, 100 μm. **(B)** DCN-specific *Cbln1* cKO caused dramatic commissural axon growth defects *in vivo*. Commissural axons were marked by Robo3 by immunostaining in spinal cord open-books at E10.5. The lengths of commissural axons are much shorter and the numbers of commissural axons are much fewer in *Cbln1* cKO spinal cords compared with their littermate controls. Notice that these differences are more obvious in posterior ends of spinal cords. A, anterior; P, posterior. Scale bar, 200 μm. **(C, D)** Quantification of commissural axon numbers and lengths in (**B**). The spinal cords were divided to bins (500 μm) along the anterior-posterior (A→P) direction and the lengths and numbers of commissural axons in each bin were quantified. All data are mean ± SEM: *Cbln1^fl/fl^* (*n* = 9 embryos) *vs Wnt1-Cre^+/-^,Cbln1^fl/fl^* (*n* = 12 embryos); *\*p* = 0.014 for Bin 6 in (**C**); *\*p* = 0.015 for Bin 3 in (**D**); *\*p* = 0.049 for Bin 4 in (**D**); *\*p* = 0.041 for Bin 5 in (**D**); by unpaired Student’s *t* test. **(E)** Unilateral DCN-specific overexpression of *cCbln1* by *in ovo* electroporation of pMath1-eGFP-IRES-cCbln1 enhanced commissural axon growth in chick neural tubes. Lhx9 marks dI1 DCNs and eGFP marks electroporated DCNs and their axons. The arrows point commissural axon terminals. Shown are the representative images from 10 chick embryos with pMath1-eGFP-IRES-cCbln1 and 8 embryos with control plasmid. **(F, G)** Quantification of commissural axon length and Lhx9^+^ neuron numbers in (**E**). All data are represented as box and whisker plots: for (**F**), pMath1-eGFP-IRES-cCbln1 (*n* = 35 sections) vs pMath1-eGFP (*n* = 31 sections), \*\*\**\*p* = 5.56E-12; for (**G**), pMath1-eGFP-IRES-cCbln1 (*n* = 37 sections) vs pMath1-eGFP (*n* = 29 sections), *p* = 0.32, ns, not significant; by unpaired Student’s *t* test.

To further test whether Cbln1 is sufficient to stimulate commissural axon growth *in vivo*, we used a model of chick neural tube. Chick *Cbln1* (*cCbln1*) is expressed in the dorsal commissural neurons (DCN) of developing chick neural tube, as is the case with mouse *Cbln1*, but is not detected in the floor plate of chick embryonic spinal cord (*Figure 2—figure supplement 1E*). We made a DCN-specific overexpression plasmid, pMath1-eGFP-IRES-MCS, by modifying a DCN-specific knockdown plasmid pMath1-eGFP-miRNA ***(Wilson and Stoeckli 2013)***. Unilateral DCN-specific overexpression of *cCbln1* by *in ovo* electroporation of pMath1-eGFP-IRES-cCbln1 enhanced chick commissural axon growth compared with control plasmid without changing commissural neuron numbers (*Figure 2E-G*). These data suggest that Cbln1 is sufficient to stimulate commissural axon growth.

Next, we continued to elucidate the mechanisms for the cell-autonomous functions of Cbln1. We hypothesized that Cbln1 was secreted from the dorsal commissural neurons (DCN) and then acted to stimulate commissural axon growth in an autocrine manner. To test this, we cultured DCN explants from E10.5 (a stage when most commissural axons have not projected to the midline yet and are called pre-crossing axons) mouse spinal cords and used Tag1 immunostaining to visualize pre-crossing commissural axons. Tag1 has been widely used as a marker for pre-crossing commissural axons ***(Chen et al. 2008; Colak et al. 2013)***. Compared with control embryonic DCN explants, the commissural axon growth of Wnt1-Cre-mediated *Cbln1* cKO DCNs was significantly inhibited, indicated by decreased axon numbers and reduced axon lengths (*Figure 3A-C*), which is consistent with *in vivo* results for DCN-specific *Cbln1* cKO (*Figure 2B-D*). These axon growth defects were efficiently rescued by adding a recombinant human Cbln1 protein (rhCbln1) to the cultures (*Figure 3A-C*). These data suggest that the cell-autonomous Cbln1 regulates commissural axon growth in an autocrine manner.

**Figure 3.**
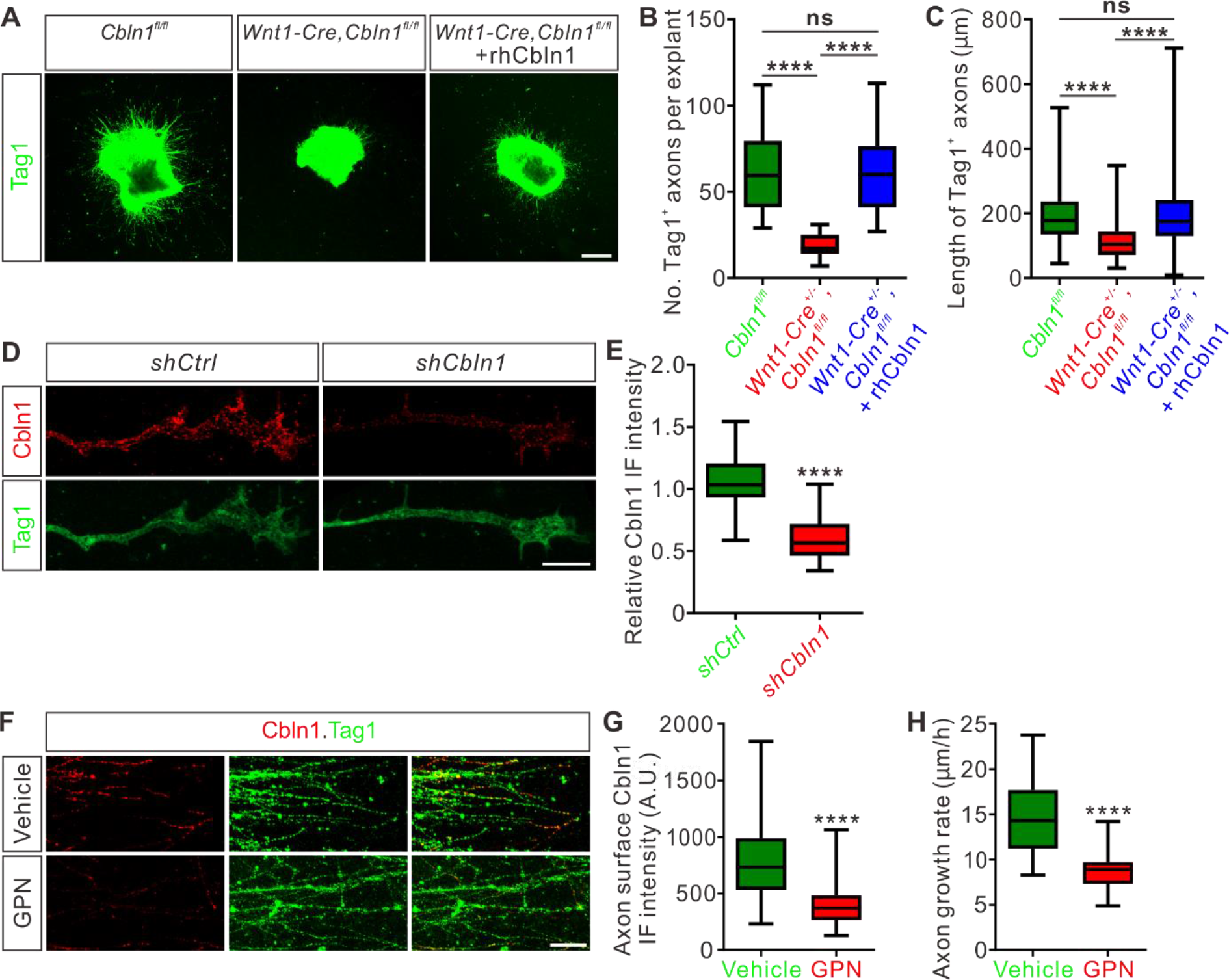
Cbln1 is secreted from commissural axon growth cones and stimulates commissural axon growth in an autocrine manner. **(A)** Extrinsic Cbln1 could rescue commissural axon growth defects caused by cell-autonomous ablation of *Cbln1* in the dorsal commissural neurons (DCN). DCN explants dissected from E10.5 mouse embryos were cultured *in vitro* and commissural axon length was monitored by immunostaining of Tag1, a commissural axon marker. Compared with *Cbln1^fl/fl^*, DCN explants of *Wnt1-Cre^+/-^,Cbln1^fl/fl^* embryos showed significant commissural axon growth defects. These defects were rescued by adding the recombinant human Cbln1 protein (rhCbln1, 500 ng/ml) to the cultures. Scale bar, 200 μm. **(B, C)** Quantification of Tag1^+^ commissural axon numbers and lengths in (**A**). Data are represented as box and whisker plots. For (**B**), \*\*\**\*p* = 1.69E-06, *Cbln1^fl/fl^* (*n* = 14 explants) vs *Wnt1-Cre^+/-^,Cbln1^fl/fl^* (*n* = 14 explants); \*\*\**\*p* = 6.23E-07, *Wnt1-Cre^+/-^,Cbln1^fl/fl^* vs *Wnt1-Cre^+/-^,Cbln1^fl/fl^* + rhCbln1 (n = 16 explants); ns, not significant (*p* = 0.99), *Cbln1^fl/fl^* vs *Wnt1-Cre^+/-^,Cbln1^fl/fl^* + rhCbln1. For (**C**), \*\*\**\*p* = 6.61E-11, *Cbln1^fl/fl^* (*n* = 876 axons) vs *Wnt1-Cre^+/-^,Cbln1^fl/fl^* (*n* = 274 axons); \*\*\**\*p* = 6.61E-11, *Wnt1-Cre^+/-^,Cbln1^fl/fl^* vs *Wnt1-Cre^+/-^,Cbln1^fl/fl^* + rhCbln1 (*n* = 1013 axons); ns, not significant (*p* = 0.91), *Cbln1^fl/fl^* vs *Wnt1-Cre^+/-^,Cbln1^fl/fl^* + rhCbln1. By one-way analysis of variance (ANOVA) followed by Tukey’s multiple comparison test. **(D)** Robust Cbln1 IF signals were detected in commissural axons and growth cones. Dissociated DCN neurons from E11 mouse embryos were cultured *in vitro* and Cbln1 IF signals were imaged after lentiviral shRNA infection. Loss of Cbln1 IF signals after *shCbln1* infection indicated the specificity of Cbln1 IF signals in commissural axons and growth cones. Scale bar, 10 μm. **(E)** Quantification of axonal Cbln1 IF signals in (**D**). Data are represented as box and whisker plots: *shCtrl* (*n* = 55 axons) vs *shCbln1* (*n* = 68 axons), \*\*\**\*p* = 5.65E-31, by unpaired Student’s *t* test. **(F)** Cbln1 is exocytosed from commissural axons via lysosomes. Robust Cbln1 IF signals were detected on the commissural axon surface of cultured DCN explants and were eliminated after blocking exocytosis with GPN treatment for 10 min. Scale bar, 50 μm. **(G)** Quantification of axon surface Cbln1 IF signals in (**F**). Data are represented as box and whisker plots: Vehicle (*n* = 140 axons) vs GPN (*n* = 126 axons), \*\*\**\*p* = 1.94E-26, by unpaired Student’s *t* test. **(H)** Blocking Cbln1 exocytosis in CA axons with GPN for 7 h inhibited CA axon growth. Data are represented as box and whisker plots: Vehicle (*n* = 75 axons) vs GPN (*n* = 51 axons), \*\*\**\*p* = 2.15E-20, by unpaired Student’s *t* test.

We next asked whether Cbln1-induced axonal growth works locally in commissural axons and growth cones. Immunofluorescence of DCN neuron culture using a Cbln1 antibody detected robust Cbln1 IF signals in commissural axons and growth cones (*Figure 3D*). To confirm the specificity of these axonal Cbln1 IF signals, we generated lentiviral *shCbln1* which led to dramatic knockdown of *Cbln1* in cultured neurons (*Figure 3—figure supplement 1A*). The Cbln1 IF signals in commissural axons and growth cones were largely lost after knockdown of Cbln1 (*Figure 3D,E*), indicating that Cbln1 is present in commissural axons and growth cones.

To test whether Cbln1 is exocytosed from axons, we applied glycyl-L-phenylalanine 2-naphthylamide (GPN) to the DCN cultures. GPN can be specifically cleaved by cathepsin C, which leads to targeted disruption of the lysosomal membrane ***(Padamsey et al. 2017; Ibata et al. 2019)***. Treatment of DCN cultures with GPN, followed by an IF protocol to detect surface Cbln1 by leaving out the permeabilization steps, showed a loss of Cbln1 IF signals on the commissural axon surface (*Figure 3F,G*), suggesting that Cbln1 is released from lysosomes in commissural axons and growth cones. Blocking Cbln1 secretion by GPN inhibited commissural axon growth (*Figure 3H*), further supporting a model that Cbln1 is released from and works back on commissural axon and growth cones to stimulate axon growth.

### Non-cell-autonomous Cbln1 from the floor plate regulates commissural axon guidance

The facts that the secreted Cbln1 works extrinsically and that it is expressed in the floor plate (FP) during commissural axon growth to the midline suggest that Cbln1 from FP might regulate commissural axon guidance. To test this idea, we first prepared COS7 cell lines stably expressing mouse Cbln1. High levels of Cbln1 were detected in the culture media, indicating the overexpressed Cbln1 was secreted from COS7 cells (*Figure 4—figure supplement 1A*). We then co-cultured the dorsal spinal cord explants from E11 mouse embryos with COS7 cell aggregates expressing Cbln1 tagged with FLAG and GFP or GFP alone in collagen gels (*Figure 4A*). The dorsal spinal cord explants growing with Cbln1-expressing COS7 cell aggregates had significantly longer axons than the control (*Figure 4A,B*). More importantly, the growth of commissural axons was attracted toward Cbln1-expressing COS7 cell aggregates, indicated by the higher axon number ratios (Proximal/Distal) compared with the control cell aggregates expressing GFP alone (*Figure 4A,C*). These results suggest that the non-cell-autonomous Cbln1 functions as an attractive axon guidance molecule.

**Figure 4.**
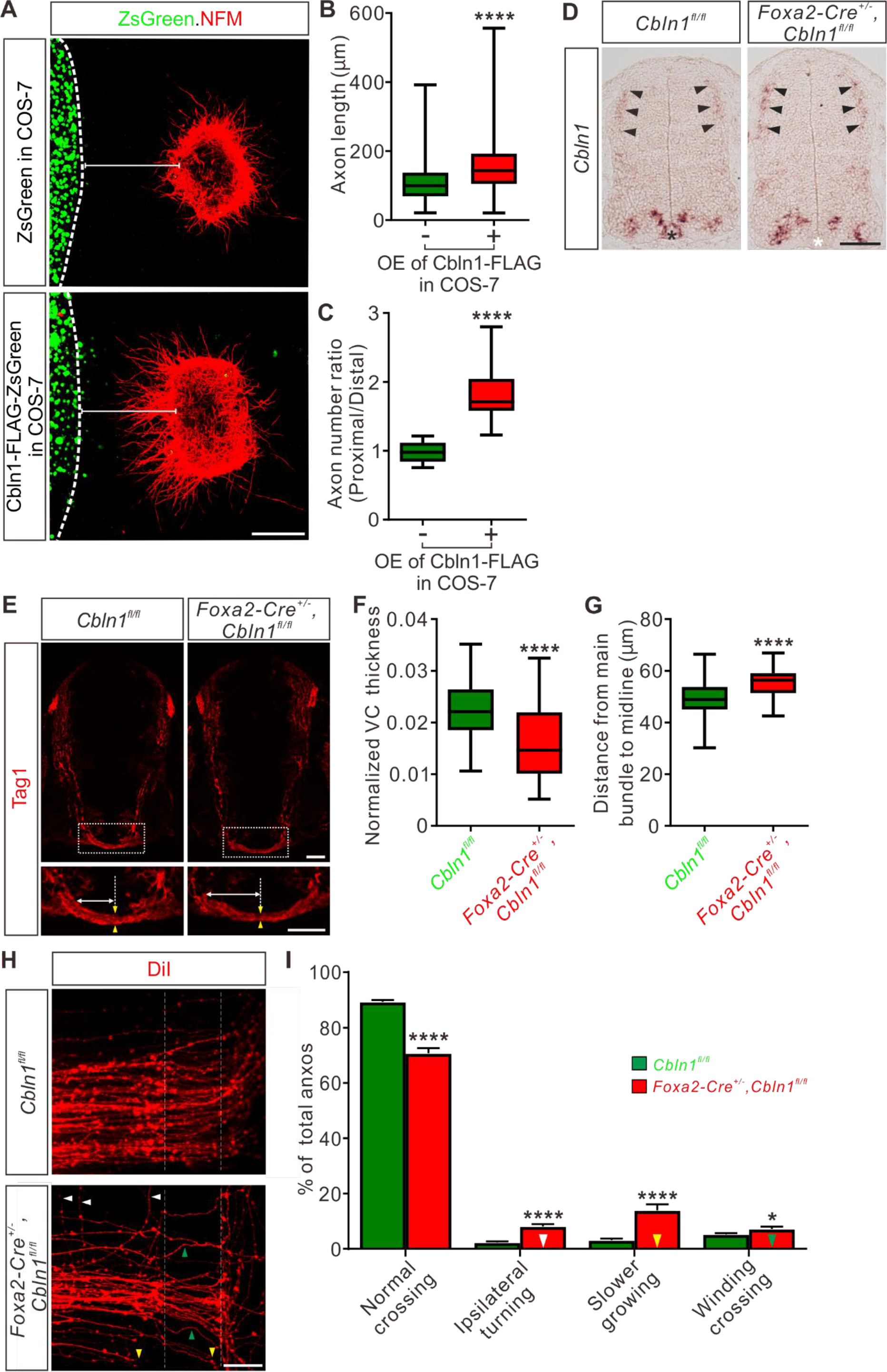
Non-cell-autonomous Cbln1 from the floor plate regulates commissural axon guidance. **(A)** Co-culture of DCN explants from E11 mouse spinal cords with COS7 cell aggregates expressing Cbln1-FLAG with ZsGreen or ZsGreen alone. Commissural axons were visualized with NFM immunostaining. Cbln1 expression attracted commissural axon turning toward cell aggregates and also enhanced axon growth. Scale bar, 200 μm. **(B, C)** Quantification of commissural axon growth and turning in (**A**) by measuring the axon length (**B**) and the axon number ratio (proximal/distal) (**C**). All data are represented as box and whisker plots: for (**B**), Ctrl (*n* = 1336 axons) vs OE (*n* = 1051 axons), \*\*\**\*p* = 1.93E-70; for (**C**), Ctrl (*n* = 16 explants) vs OE (*n* = 14 explants), \*\*\**\*p* = 1.97E-08; by unpaired Student’s *t* test. **(D)** Specific ablation of *Cbln1* in the floor plate of *Foxa2-Cre^+/-^,Cbln1^fl/fl^* cKO mouse embryos was confirmed by *in situ* hybridization of E11.5 spinal cord sections. Expression of *Cbln1* in the floor plate was completely lost in the cKO spinal cord (white asterisk) compared with the control embryos (black asterisk). Black arrowheads indicate the unchanged *Cbln1* expression in DCNs of both genotypes. Scale bar, 100 μm. **(E)** The axon guidance defects of pre-crossing commissural axons were observed by Tag1 immunostaining in floor plate-specific *Cbln1* cKO and control embryos at E11.5. Higher magnification views of the floor plate region in the white dotted boxes are also shown (bottom). The pair of yellow arrowheads denotes the thickness of the ventral commissure (VC). The double-arrowed line measures the distance between the point of intersection (of the main pre-crossing commissural axon bundle with the ventral edge of spinal cord) and the midline (indicated by the dotted line). Scale bars, 50 μm. **(F, G)** Quantification of the VC thickness and the distance from the main bundle intersection point to the midline. The VC thickness was normalized to the height (dorsal to ventral) of spinal cord. All data are represented as box and whisker plots: *Cbln1^fl/fl^* (*n* = 62 sections) vs *Foxa2-Cre^+/-^,Cbln1^fl/fl^* (*n* = 60 sections), \*\*\**\*p* = 1.69E-06 for (**F**), \*\*\**\*p* = 1.07E-07 for (**G**), by unpaired Student’s *t* test. **(H)** DiI labeling of E11.5 spinal cord open-books traced commissural axon guidance behaviors during midline crossing. The region between two white dotted lines indicates the floor plate. The white, yellow and green arrowheads indicate the commissural axons with aberrant behaviors such as ipsilateral turning, slower growing or winding crossing, respectively. Scale bar, 50 μm. **(I)** Quantification of the percentages of commissural axons with different guidance behaviors. All data are mean ± SEM. and represented as histogram: *Cbln1^fl/fl^* (*n* = 45 DiI injections) vs *Foxa2-Cre^+/-^,Cbln1^fl/fl^* (*n* = 32 DiI injections), \*\*\**\*p* = 3.36E-17 for normal crossing, \*\*\**\*p* = 8.86E-10 for ipsilateral turning, \*\*\**\*p* = 6.14E-07 for slower growing, *\*p* = 0.033 for winding crossing, by unpaired Student’s *t* test.

To assess the *in vivo* functions of the non-cell-autonomous Cbln1, we generated floor plate-specific *Cbln1* cKO mice. We utilized *Foxa2-Cre^ERT^* line which has been used to induce Cre recombinase expression specifically in floor plate cells in response to tamoxifen (TM) treatment ***(Park et al. 2008; Hernandez-Enriquez et al. 2015)***. *Cbln1* expression was specifically ablated from the floor plate in these cKO embryos, without affecting its expression in other parts of spinal cord including the dorsal commissural neurons (DCNs) (*Figure 4D*). The neural patterning or neurogenesis was not disturbed by ablation of Cbln1 from the floor plate (FP) (*Figure 4—figure supplement 1B-E*). However, examination of commissural axon trajectories using Tag1 immunostaining in E11.5 spinal cords revealed significant axon guidance defects in the midline and ventral spinal cord. First, the thickness of the ventral commissure (VC) was significantly reduced in the FP-specific *Cbln1* cKO embryos compared with their littermate controls (*Figure 4E,F*). Second, the intersection of the main commissural axon bundle with the ventral commissural funiculus was shifted laterally in the FP-specific *Cbln1* cKO embryos compared with their littermate controls (*Figure 4E*). The distances between the point of intersection and the midline were quantified, showing a significant increase in the FP-specific *Cbln1* cKO embryos (*Figure 4G*). These phenotypes were also evident by NFM immunostaining (*Figure 4—figure supplement 1F-H*). These axon guidance defects suggest that Cbln1 from the floor plate indeed works as an axon guidance cue in the developing spinal cord.

To observe more clearly the commissural axon guidance behaviors in the FP-specific *Cbln1* cKO embryos, we performed DiI labeling of DCNs in the open-book spinal cords at E11.5. As shown in *Figure 4H,I*, there was a significant decrease of the number of normal crossing commissural axons in the *Cbln1* cKO. Meanwhile, the numbers of commissural axons showing guidance defects such as ipsilateral turning, slower growing or winding crossing were significantly increased in *Cbln1* cKO embryos compared with their littermate controls (*Figure 4H,I*).

All these data support the idea that the non-cell-autonomous Cbln1 derived from the floor plate works as an axon guidance cue for commissural axons in the developing spinal cord.

### Nrxn2 is expressed in the developing dorsal commissural neurons and axons, and mediates Cbln1-induced axon growth and guidance as its receptor

Previously Cbln1 was shown to work as a synaptic organizer for the cerebellar excitatory PF-PC (PF, parallel fibers; PC, Purkinje cells) synapses by binding to its presynaptic receptor, neurexin (Nrxn) and its postsynaptic receptor, glutamate receptor delta 2 (GluD2) ***(Matsuda et al. 2010; Uemura et al. 2010)***. In addition, trans-synaptic signaling through Nrxn-Cbln-GluD1 has also been shown to mediate the inhibitory synapse formation in cortical neurons ***(Yasumura et al. 2012; Fossati et al. 2019)***. Here we wondered whether Nrxn, GluD1, and/or GluD2 work as receptors for Cbln1 to mediate its regulation of commissural axon growth and guidance in the developing spinal cord. To test this, we first checked if Nrxn, GluD1, and GluD2 are expressed in developing dorsal commissural neurons (DCNs) or not. *GluD1* or *GluD2* mRNA was not detected in E11.5 spinal cords (*Figure 5—figure supplement 1A*). There are three *Nrxn* genes in the mammalian genome, each of them encoding two major protein isoforms, α-neurexin and β-neurexin ***(Reissner et al. 2013)***. Although *Nrxn1*, *2* and *3* mRNAs were all detected in E11.5 spinal cords, only *Nrxn2* mRNA was found to be expressed in DCNs (*Figure 5A*). We further detected both *Nrxn2α* and *Nrxn2β* mRNAs in DCNs using the isoform-specific probes (*Figure 5B*). Immunostaining using an Nrxn2 antibody detected robust Nrxn2 IF signals in commissural axons and growth cones (*Figure 5C,D*), making it possible that Nrxn2 works as the receptor for Cbln1 in developing commissural axons. Indeed, commissural axon lengths were significantly decreased after knocking down either pan-Nrxn2 or Nrxn2α, Nrxn2β separately (*Figure 5—figure supplement 1B-D*, and *Figure 5E,F*), suggesting that Nrxn2 mediates cell-autonomous-Cbln1-induced commissural axon growth. We continued to test whether Nrxn2 also mediates the non-cell-autonomous function of Cbln1 to attract commissural axon turning. As shown in *Figure 5G*, Cbln1-expressing cell aggregates failed to attract commissural axons of DCN neurons which were knocked down of either pan-Nrxn2 or Nrxn2α, Nrxn2β separately. These data suggest that Nrxn2 (both Nrxn2α and Nrxn2β) works as the receptor for Cbln1 to mediate its cell-autonomous function in axon growth and non-cell-autonomous function in axon guidance of commissural neurons in the developing spinal cord.

**Figure 5.**
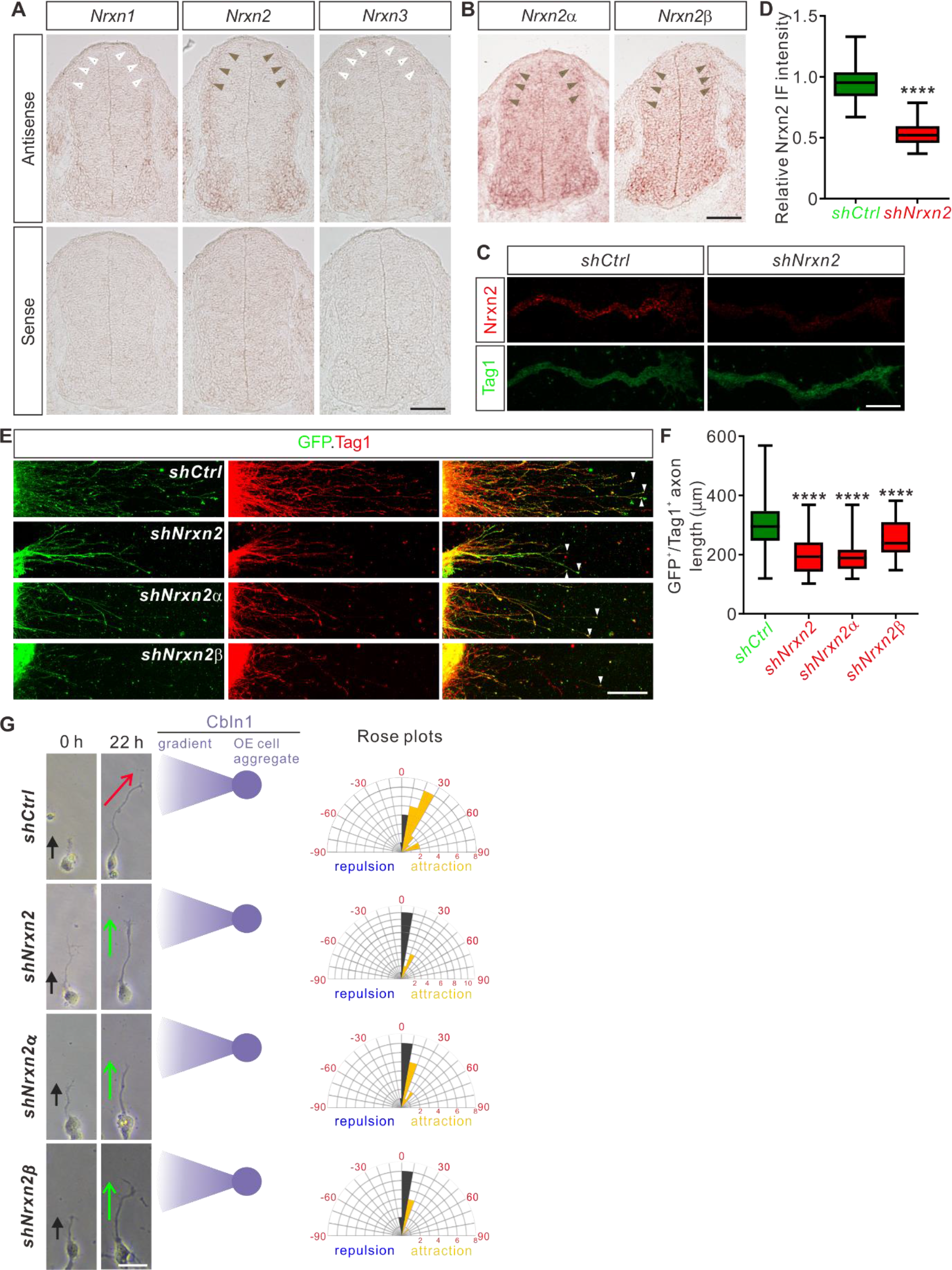
Nrxn2 mediates Cbln1-induced commissural axon growth and guidance as its receptor. **(A, B)** *Nrxn2*, *Nrxn2α* and *Nrxn2β* mRNAs were detected in E11.5 spinal cord cross-sections by *in situ* hybridization. Brown and white arrowheads indicate the expression (*Nrxn2* in **A**, *Nrxn2α* and *Nrxn2β* in **B**) or absence (*Nrxn1* and *Nrxn3* in **A**) of the corresponding mRNAs in DCNs, respectively. Scale bars, 100 μm. **(C)** Robust Nrxn2 IF signals were detected in the commissural axons and growth cones. Dissociated DCN neurons from E11 mouse embryos were cultured *in vitro* and Nrxn2 IF signals were imaged after lentiviral shRNA infection. Loss of Nrxn2 IF signals after *shNrxn2* infection indicated the specificity of Nrxn2 IF signals in the commissural axons and growth cones. Scale bar, 10 μm. **(D)** Quantification of axonal Nrxn2 IF signals in (**C**). Data are represented as box and whisker plots: *shCtrl* (*n* = 69 axons) vs *shNrxn2* (*n* = 65 axons), \*\*\**\*p* = 3.75E-39, by unpaired Student’s *t* test. **(E)** Knockdown of Nrxn2, Nrxn2α or Nrxn2β in DCNs inhibited commissural axon growth. Axons of DCN neurons which were infected by shRNA against *Nrxn2* were marked by both GFP reporter and Tag1 IF. White arrowheads indicate the axon terminals. Scale bar, 100 μm. **(F)** Lengths of GFP^+^/Tag1^+^ commissural axons in (**E**) were measured and analyzed. Data are represented as box and whisker plots: *shCtrl* (*n* = 63 axons) vs *shNrxn2* (*n* = 53 axons), \*\*\**\*p* = 1.41E-15; *shCtrl* vs *shNrxn2α* (*n* = 48 axons), \*\*\**\*p* = 4.30E-18; *shCtrl* vs *shNrxn2β* (*n* = 48 axons), \*\*\**\*p* = 6.27E-05; by one-way analysis of variance (ANOVA) followed by Tukey’s multiple comparison test. **(G)** Knockdown of Nrxn2, Nrxn2α or Nrxn2β in DCNs disturbed commissural axon turning toward Cbln1-expressing cell aggregates. Dissociated DCN neurons from E11 mouse spinal cords were infected with shRNAs, and co-cultured with COS7 cell aggregates expressing Cbln1. Commissural axons were imaged at two time points (0 and 22 h). Commissural axon turning angles toward the Cbln1-OE cell aggregates and gradients were measured between the colored arrow (red for *shCtrl* and green for *shNrxn2s*) at 22 h and the black arrow at 0 h. Rose plots of axon turning angles are shown to the right for each condition. Angles were clustered in bins of 10°, and the number of axons per bin is represented by the radius of each segment. Orange bins indicate attraction, and blue bins indicate repulsion. Scale bar, 20 μm.

In summary, these data and findings support the following working model for Cbln1 in the developing spinal cord (*Figure 5—figure supplement 1E,F*). In the pre-crossing commissural axons, Cbln1 is expressed cell-autonomously by the dorsal commissural neurons (DCN) and axons.

Commissural axon growth cone-secreted Cbln1 binds to Nrxn2 receptors in commissural axons and growth cones to stimulate commissural axon growth in an autocrine manner (*Figure 5—figure supplement 1E*). In the DCN-specific *Cbln1* cKO embryos, commissural axon growth is reduced compared with their littermate controls (*Figure 5—figure supplement 1E*). When commissural axons approach the midline, the floor plate-derived Cbln1 attracts commissural axons to the midline which is also mediate by Nrxn2 receptors (*Figure 5—figure supplement 1F*). In the floor plate-specific *Cbln1* cKO embryos, commissural axon guidance in the midline crossing is impaired, resulting in a U-shaped and thinner ventral commissure compared with the V-shaped and thick ventral commissures in the littermate control embryos (*Figure 5—figure supplement 1F*).

### Cell-autonomous Cbln1 from cerebellar granular cells is required for parallel fiber growth

We wondered whether the function of Cbln1 to regulate axon development is a general mechanism which also works in other brain regions during development. The studies on Cbln1 so far have been focused on its functions as a synaptic organizer in cerebellum. Whether Cbln1 is expressed and exerts its functions at earlier stages of cerebellar development remains unexplored. We first checked expression of Cbln1 in earlier cerebellar development. As shown in *Figure 6A*, high and specific *Cbln1* expression was detected in the P4-P8 cerebellar granule cells in the inner granule layer (IGL) by *in situ* hybridization. Immunofluorescence using a Cbln1 antibody showed that Cbln1 protein is enriched in the molecular layer (ML) of cerebellum (*Figure 6B*), suggesting that Cbln1 protein is expressed and secreted by cerebellar granule cell (GC) axons.

**Figure 6.**
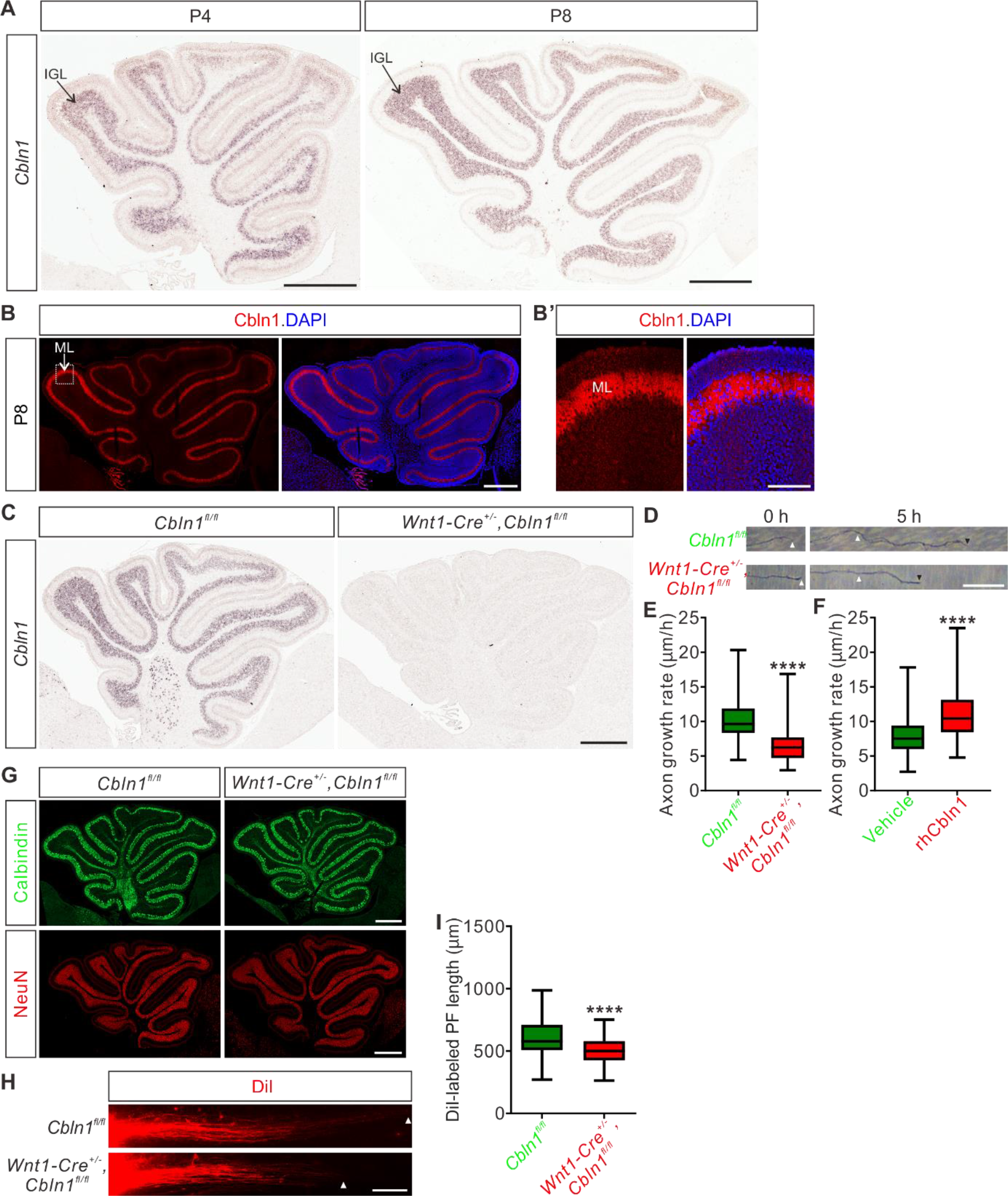
Cell-autonomous Cbln1 is required for cerebellar granule cell axon growth. **(A)** *In situ* hybridization of *Cbln1* in cerebella during P4 and P8. *Cbln1* mRNA is specifically and highly expressed in granule cells, esp. in the inner granule layer (IGL). Scale bars, 500 μm. **(B)** High level of Cbln1 protein is detected in the molecular layer (ML) of P8 cerebellum, which is expressed and secreted by granule cell (GC) axons. Higher magnification of the boxed area is shown in (**B’**). Scale bars, 500 μm (**B**) and 100 μm (**B’**). **(C)** Ablation of *Cbln1* expression in *Cbln1* cKO mouse cerebella. *In situ* hybridization of *Cbln1* in P8 cerebellum confirmed the ablation of *Cbln1* from IGL. Scale bar, 500 μm. **(D, E)** Cell-autonomous Cbln1 is required for GC axon growth. P6 GC neurons were dissected and cultured *in vitro*. GC axons were imaged at two time points (0 and 5 h). The growth rate of GC axons from *Cbln1* cKO cerebella was significantly slower than that of control. Quantification data are represented as box and whisker plots (**E**): *Cbln1^fl/fl^* (*n* = 298 axons) vs *Wnt1-Cre^+/-^,Cbln1^fl/fl^* (*n* = 247 axons); \*\*\**\*p* = 4.05E-52; by unpaired Student’s *t* test. Scale bar, 20 μm. **(F)** Extrinsic Cbln1 could stimulate GC axon growth. WT P6 GC neurons were cultured *in vitro* and rhCbln1 (500 ng/ml) was added to the culture. Compared with the vehicle control, rhCbln1 significantly enhanced GC axon growth. Data are represented as box and whisker plots: Vehicle (*n* = 342 axons) vs rhCbln1 (*n* = 299 axons); \*\*\**\*p* = 5.58E-32; by unpaired Student’s *t* test. **(G)** Neurogenesis is not disturbed in the *Cbln1* cKO cerebellum at P8. Immunostainings of the Purkinje cell marker Calbindin and the granule cell marker NeuN showed no difference between *Cbln1* cKO and control cerebella, suggesting that the neurogenesis of PCs and GCs in cerebellum is not affected. Scale bars, 500 μm. **(H, I)** Lengths of parallel fibers labeled by DiI were significantly decreased in *Cbln1* cKO mice at P6. The white arrowheads indicate the terminals of DiI-labeled PFs. Quantification of PF lengths is shown as box and whisker plots (**I**): \*\*\**\*p* = 1.09E-13; *n* = 190 axons for *Cbln1^fl/fl^* mice, *n* = 132 axons for *Wnt1-Cre^+/-^,Cbln1^fl/fl^* mice; by unpaired Student’s t-test. Scale bar, 100 μm.

Next we tested the possible roles of Cbln1 in earlier cerebellar development. We generated *Cbln1* cKO in cerebellum using the *Wnt1-cre* line ***(Danielian et al. 1998; Cerrato et al. 2018)***, which resulted in the efficient knockout of *Cbln1* from GCs (*Figure 6C*). Axon growth rates of *Cbln1*-deficient GCs *in vitro* were significantly decreased compared with control neurons (*Figure 6D,E*), suggesting that the cell-autonomous Cbln1 is required for GC axon growth. Similar to Cbln1 on commissural axons, extrinsic application of the recombinant hCbln1 (rhCbln1) protein to the GC axons stimulated their growth (*Figure 6F*), supporting a similar model as in the developing spinal cord that Cbln1 secreted from cerebellar GC axons works back to stimulate GC axon growth in the developing cerebellum. Detection of *Nrxn1, 2,* and *3* expression in GCs at IGL (*Figure 6—figure supplement 1A,B*) implied that neurexins would mediate the autocrine function of Cbln1 to stimulate GC axon growth in the developing cerebellum as in the spinal cord.

We continued to carefully examine the *Cbln1* cKO cerebella. Immunostaining of the Purkinje cell (PC) marker Calbindin and the granule cell (GC) marker NeuN showed no difference between *Cbln1* cKO and control cerebella at P8 (*Figure 6G*), suggesting that the neurogenesis of PC and GC in the cerebellum is not impaired. To investigate whether the *in vitro* regulation of GC axon growth by Cbln1 was recapitulated *in vivo*, we examined parallel fiber (PF) development in *Cbln1* cKO mice by DiI labeling. Compared with control mice, the DiI-labeled parallel fiber lengths in *Cbln1* cKO mouse pups at P6 were significantly decreased (*Figure 6H,I*), indicating that the parallel fiber growth was impaired in *Cbln1* cKO cerebella.

All these data suggested that cell-autonomous Cbln1 from granule cells is required for parallel fiber growth in the developing cerebellum, just as cell-autonomous Cbln1 from commissural axons stimulates CA axon growth in the developing spinal cord.

### Non-cell-autonomous Cbln1 regulates axon guidance of retinal ganglion cells in the optic chiasm

The regulation of commissural axon guidance during midline crossing by the non-cell-autonomous Cbln1 from floor plate of the developing spinal cord inspired us to further test whether Cbln1 regulates axon guidance in other brain midline models. The optic chiasm (OC) is where retinal ganglion cell (RGC) axons from each eye cross the midline. The ipsi- and contra-lateral axon organization of RGC axons in OC is critical for binocular vision ***(Mason and Slavi 2020)***. We wanted to check whether Cbln1 contributed to axon guidance signaling in OC. RGCs do not express *Cbln1* (*Figure 7A*). However, both *Cbln1* mRNA and Cbln1 protein were detected in the ventral diencephalon at the floor of the third ventricle which is adjacent to OC (*Figure 7B,C*), implying that Cbln1 has the right location to exert effects on OC. *In vitro* co-culture of retinal explants with COS7 cell aggregates expressing Cbln1 showed that Cbln1 is sufficient to attract RGC axon turning (*Figure 7D*).

**Figure 7.**
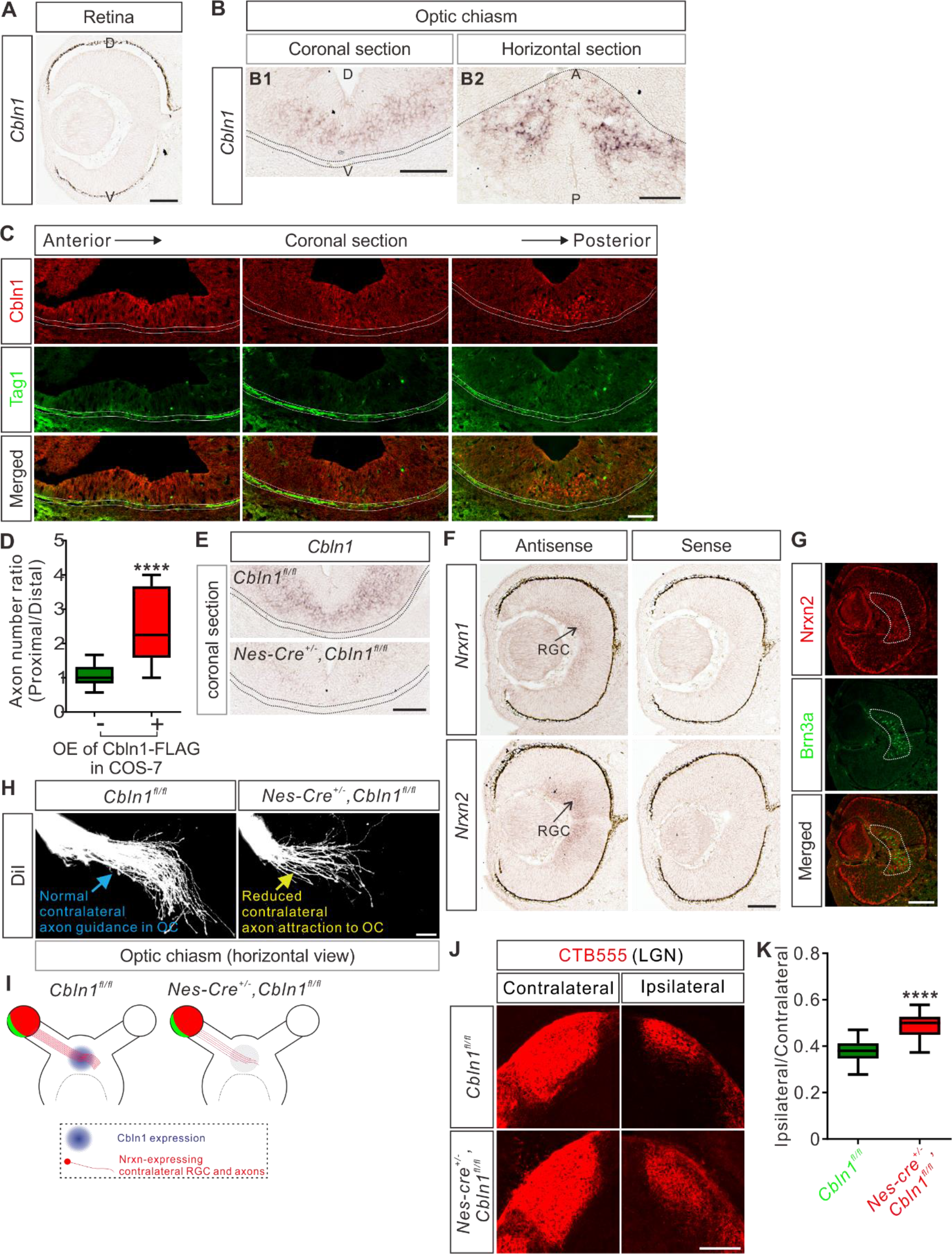
Non-cell-autonomous Cbln1 regulates RGC axon guidance in optic chiasm. **(A)** *In situ* hybridization of *Cbln1* in the developing retina. *Cbln1* mRNA expression was not detected in the retina of E13 mouse embryos. D, dorsal; V, ventral. Scale bar, 100 μm. **(B)** *In situ* hybridization of *Cbln1* in E13 mouse brain sections (coronal section for **B1**; horizontal section for **B2**). *Cbln1* mRNA expression was detected in the floor of the third ventricle, adjacent to the optic chiasm. The dotted lines indicate the boundary of the optic chiasm. D, dorsal; V, ventral; A, anterior; P, posterior. Scale bars, 100 μm. **(C)** Immunostaining of Cbln1 in coronal brain sections from E13 mouse embryos. Cbln1 protein was detected in the floor of the third ventricle, adjacent to the optic chiasm. Tag1-marked RGC axons projected to the optic chiasm are outlined by the dotted lines. Serial sections from anterior level to posterior level were shown. Scale bar, 100 μm. **(D)** Extrinsic Cbln1 attracted RGC turning *in vitro*. Retina explants from E13 mouse embryos were co-cultured with COS7 cell aggregates expressing Cbln1-FLAG with GFP or GFP alone. Quantification of RGC axon turning was performed by measuring the axon number ratio (proximal/distal), similarly as CA axon guidance assay. Data are represented as box and whisker plots: Ctrl (*n* = 17 explants) vs OE (*n* = 9 explants), \*\*\**\*p* = 5.62E-05; by unpaired Student’s *t* test. **(E)** Ablation of *Cbln1* in *Nes-Cre^+/-^,Cbln1^fl/fl^* cKO mouse embryos was confirmed by *in situ* hybridization of E13 coronal brain sections. The dotted lines indicate the boundary of the optic chiasm. Scale bar, 100 μm. **(F)** *In situ* hybridization of *Nrxn1* and *Nrxn2* in E13 retina. *Nrxn1* and *Nrxn2* mRNAs were detected in retinal ganglion cells (RGC). Scale bar, 100 μm. **(G)** Immunostaining of Nrxn2 in E13 retina. Nrxn2 is expressed only in the contralateral RGCs marked by Brn3a. Scale bar, 100 μm. **(H, I)** Axon guidance defects in the optic chiasm (OC) of *Cbln1* cKO embryos. DiI tracing of RGC axons was performed to visualize axon trajectory in OC. Compared with normal axon attraction of contralateral RGCs in OC of control embryos, *Cbln1* cKO embryos showed reduced axon attraction to OC. The phenotype is illustrated in (**I**). Scale bar, 100 μm. **(J, K)** RGC central targeting defects of *Cbln1* cKO mice. Representative images of coronal sections through the LGN (lateral geniculate nucleus) after unilateral injection of CTB-Alexa Fluor 555 at P4 in *Cbln1* cKO and control mice were shown and projections to the contralateral and ipsilateral LGN are visible (**J**). Quantification of “Ipsilateral area”/“Contralateral area” is represented as box and whisker plot (**K**): Ctrl (*n* = 51 sections) vs cKO (*n* = 53 sections), \*\*\**\*p* = 2.06E-18; by unpaired Student’s *t* test. Scale bar, 200 μm.

We next continued to explore whether Cbln1 physiologically regulates RGC axon guidance in OC at the ventral diencephalic midline. We generated *Cbln1* cKO embryos using *Nes-cre*. As show in *Figure 7E*, *Cbln1* expression in the ventral diencephalon was ablated. RGCs can be divided to ipsilateral and contralateral subgroups according to their projection laterality to the same or opposite side of the brain, respectively. The experiments checking expression of Cbln1 receptors in retina by *in situ* hybridization revealed that only *Nrxn1* and *Nrxn2* mRNA were detected in the developing retina (*Figure 7F*) while *Nrxn3*, *GluD1* or *GluD2* mRNA was not detected (*Figure 7—figure supplement 1A,B*). Nrxn2 was further found to be only expressed in the contralateral RGCs marked by Brn3a (*Figure 7G*), suggesting that Cbln1 would only work on the contralateral RGCs. Consistent with this, DiI tracing of RGC axons showed that contralateral axon attraction to OC was impaired in the *Cbln1* cKO mouse embryos compared with control embryos (*Figure 7H,I*). We further checked the targeting of optic nerves to the brain by anterograde labeling with cholera toxin subunit B (CTB) and found that the ratio of ipsilateral area to contralateral area of the retinogeniculate projections was increased in *Cbln1* cKO pups compared with control pups (*Figure 7J,K*). These data suggest that the non-cell-autonomous Cbln1 regulates contralateral RGC axon guidance in the optic chiasm.

## Discussion

Based on the *in vitro* and *in vivo* studies in mice, we have demonstrated that cell-autonomous and non-cell-autonomous Cbln1 regulates axon growth and guidance in multiple neural regions, respectively, suggesting a general role for Cbln1 in early nervous system development.

Studies on Cbln1 so far have focused on its role as the synaptic organizer in cerebellum *(Hirai et al. 2005; Matsuda et al. 2010; Uemura et al. 2010; Ito-Ishida et al. 2012; Yuzaki 2018; Ibata et al. 2019; Suzuki et al. 2020; Takeo et al. 2020)* and cortex *(Fossati et al. 2019)*. Whether Cbln1 works in earlier neuronal developmental processes prior to synapse formation or in other neural regions is not known. Here we report that Cbln1 is expressed in developing spinal cord, cerebellum, and ventral diencephalon. We also found that Cbln1 regulates axon growth and guidance in multiple neural regions. These findings suggest that Cbln1 has dynamic spatial-temporal expression and function in the nervous system. Thus, in order to distinguish the early (axon development) and late (synapse formation) roles of Cbln1, it would be critical to more precisely control the timepoint of knocking out *Cbln1*. For example, inducible *Cbln1* cKO would be necessary to explore its roles in synapse formation, in order to avoid disrupting its role in axon pathfinding.

During neural developmental stages before synapse formation, extracellular cues are required to direct axon growth and guidance ***(Stoeckli 2018)***. Most of these cues are non-cell-autonomous and secreted by sources such as the surrounding and target (intermediate or final) tissues. Here we found that non-cell-autonomous Cbln1 expressed and secreted from floor plate in the developing spinal cord and from ventral diencephalon in the developing brain works in a paracrine manner to regulate commissural axon and retinal ganglion axon guidance when they cross the midline.

In addition, we also found that cell-autonomous Cbln1 which is generated and secreted from commissural and cerebellar granule cell axons works in an autocrine manner to stimulate their own axon growth. Actually, other examples that axon-derived and remotely secreted cues regulate axon development have also been reported. The axonally secreted protein axonin-1 promotes neurite outgrowth of dorsal root ganglia (DRG) ***(Stoeckli et al. 1991)***. Wnt3a is expressed in RGCs and has the autocrine RGC axon growth-promoting activity ***(Harada et al. 2019)***. The C terminus of the ER stress-induced transcription factor CREB3L2 was found to be secreted by DRG axons to promote DRG axon growth ***(McCurdy et al. 2019)***. Recent study suggests that axonally synthesized Wnt5a is secreted and promote cerebellar granule axon growth in an autocrine manner ***(Yu et al. 2021)***.

Thus, Cbln1 shows up as an example of molecules with dual roles as both non-cell-autonomous and cell-autonomous cues to regulate axon guidance and growth, respectively.

We found that neurexins, esp. Nrxn2, mediate Cbln1 functions in axon development. Neurexins are transmembrane proteins with a large extracellular region and a small intracellular C-terminal region ***(Reissner et al. 2013)***. *Nrxns* are alternatively spliced at six sites (named as SS1 to SS6)***(Sudhof 2017)***, whereas Cbln1 only bind to SS4+ neurexins ***(Uemura et al. 2010)***. Extracellularly, α-Nrxns bind to Cbln1 via the LNS6 domain (laminin/neurexin/sex-hormone-binding globulin domain 6) which is also shared by β-Nrxns ***(Sudhof 2017)***. Intracellularly, Nrxns interact with CASK, Mints, and protein 4.1 which nucleates actin cytoskeleton to regulate synapse formation ***(Hata et al. 1996; Biederer and Südhof 2000; Biederer and Sudhof 2001; Mukherjee et al. 2008)***. It will be interesting to explore whether and how Cbln1-Nrxn signaling is mediated by intracellular CASK/Mint/p4.1-cytoskeleton pathway to regulate axon growth and guidance in the early neuronal development.

## Materials and methods

Key resources table

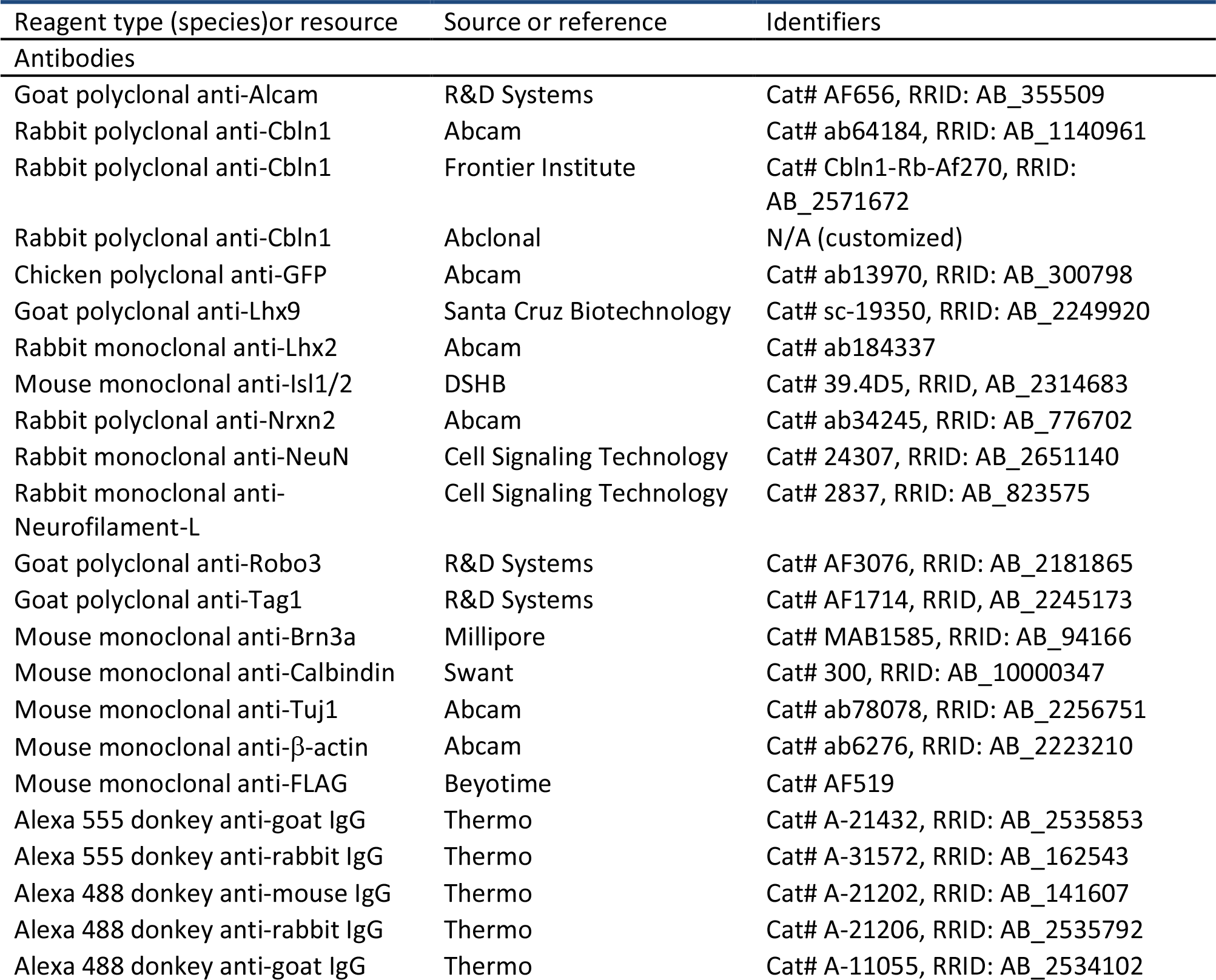
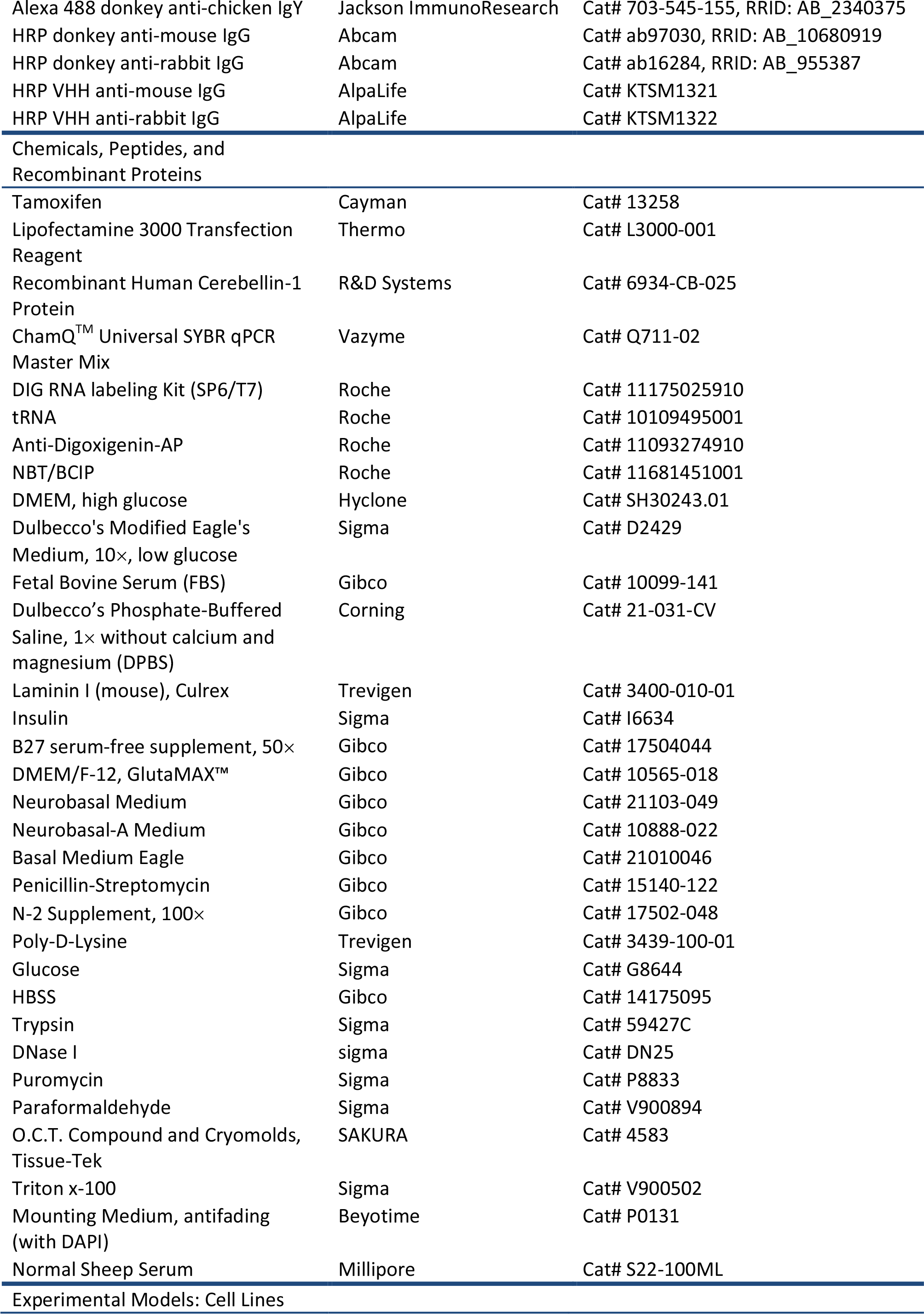
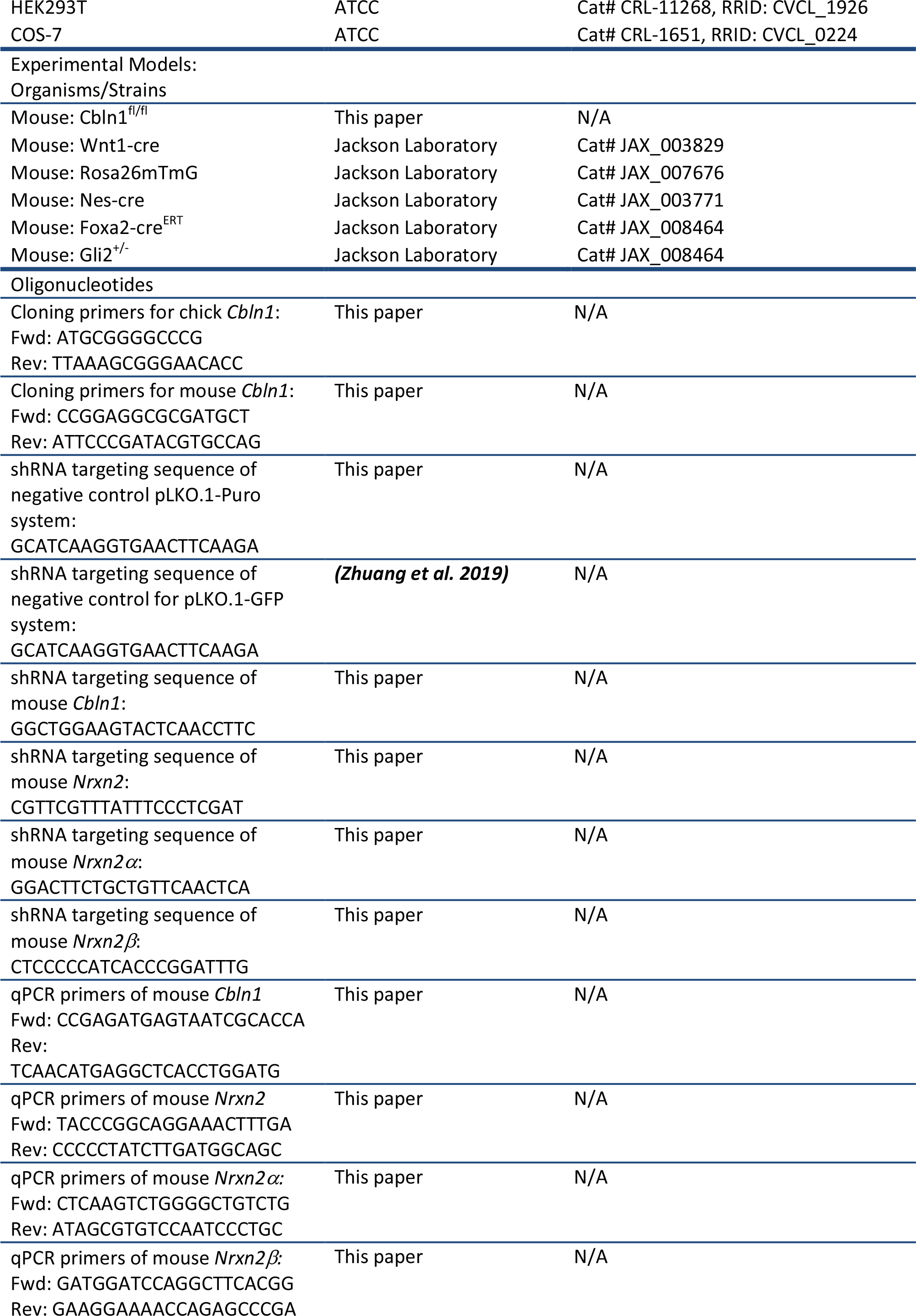
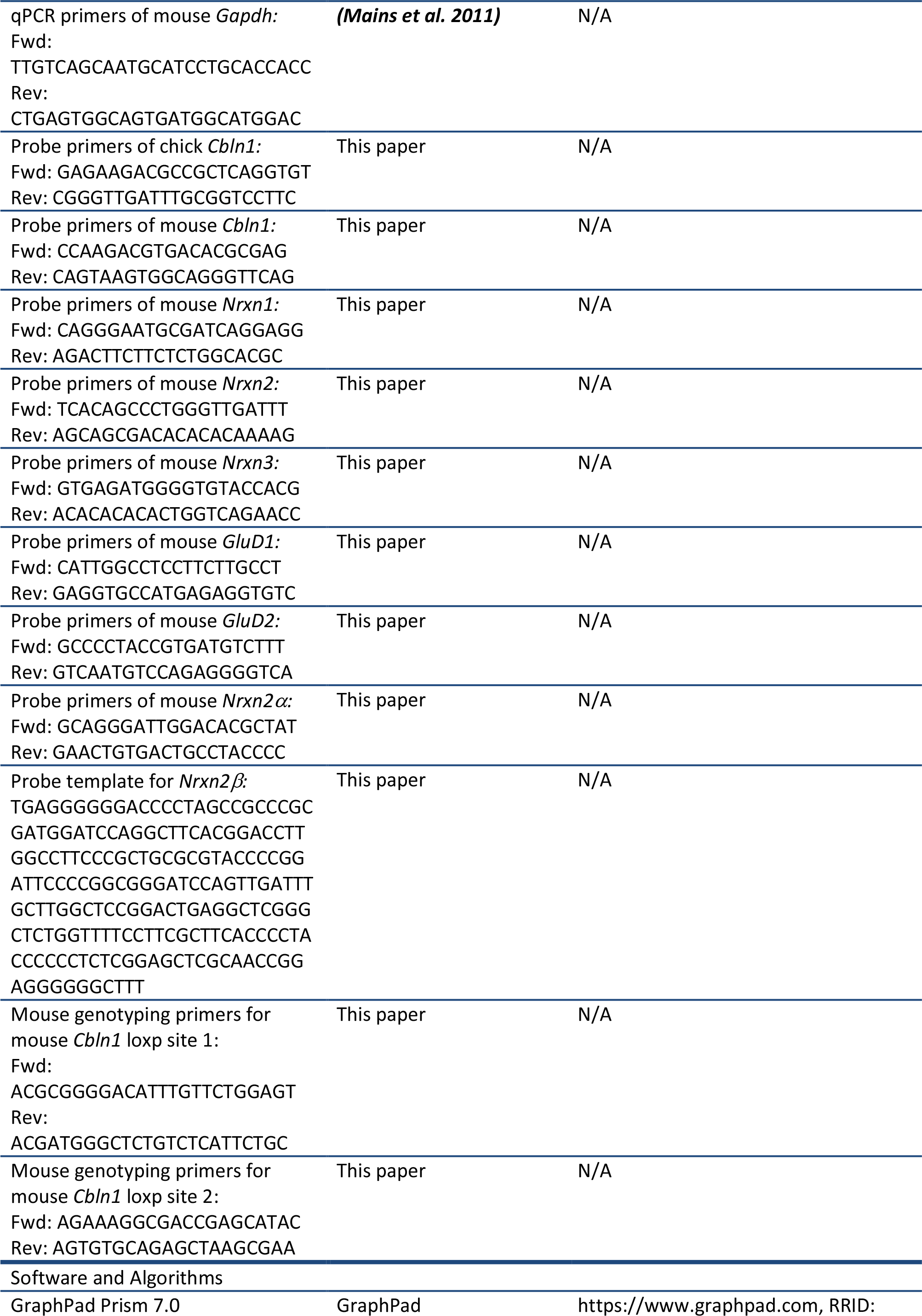
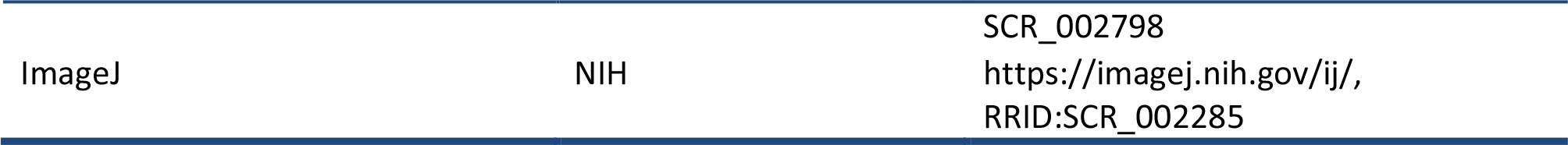

### Animals

Generation of *Cbln1* conditional knockout (cKO) mice was performed following procedures described previously ***(Zhuang et al. 2019)***, with the whole coding sequence as the targeted region (*Figure 2— figure supplement 1A*). *Cbln1^fl/+^* mice and corresponding *Cre* mice lines were used to generate *Cbln1* cKO and littermate control embryos. Genotyping primers are as following: the first *Cbln1-loxP* site, 5’-ACGCGGGGACATTTGTTCTGGAGT-3’ and 5’-ACGATGGGCTCTGTCTCATTCTGC-3’; the second *Cbln1-loxP* site, 5’-AGAAAGGCGACCGAGCATAC-3’ and 5’-AGTGTGCAGAGCTAAGCGAA-3’. *Wnt1-cre **(Danielian et al. 1998)***, *Rosa26mTmG **(Muzumdar et al. 2007)***, *Gli2^+/-^ **(Bai and Joyner 2001)***, *Foxa2-cre^ERT^ **(Park et al. 2008)***, and *Nes-cre **(Tronche et al. 1999)*** mice used in this study were described in the indicated references and their stock numbers in The Jackson Laboratory are 003829, 007676, 007922, 008464, and 003771 respectively. All mice were housed in a specific pathogen-free animal facility at the Laboratory Animal Center of Southern University of Science and Technology. All experiments using mice were carried out following animal protocols approved by the Laboratory Animal Welfare and Ethics Committee of Southern University of Science and Technology. For timed pregnancy, embryos were identified as E0.5 when a copulatory plug was observed at noon. To induce Cre activity for *Foxa2-cre^ERT^*-derived *Cbln1* cKO in floor plate, 8 mg tamoxifen (Cayman Chemical) was given orally to E8.5 pregnant mice with an animal gauge feeding needle. Fertilized chick eggs were purchased from a local supplier and chick embryos developed in an incubator (BSS 420, Grumbach) were staged using the Hamburger and Hamilton staging system. For all experiments with mice or chick, a minimum of three (up to 20) embryos or pups was analyzed for each genotype or experimental condition.

### In ovo electroporation

The chick spinal DCN-specific knockdown vector pMath1-eGFP-miRNA was a gift from Esther T. Stoeckli ***(Wilson and Stoeckli 2013)***. DCN-specific overexpression vector pMath1-eGFP-IRES-MCS was constructed by replacing the miRNA cassette with an IRES sequence plus multiple coning sites (MCS). The coding sequence of chick *Cbln1* was cloned from St. 23/24 chick spinal cord cDNA with primers 5’-ATGCGGGGCCCG-3’ and 5’-TTAAAGCGGGAACACC-3’. *In ovo* electroporation was carried out using the ECM® 830 Square Wave Electroporator (BTX) as previously described ***(Ji et al. 2009)***. Electroporation was performed at St.17 and embryos were collected and analyzed at St.23.

### Tissue and neuron culture

All tissue culture reagents were from Thermo unless otherwise specified. DCN explant and neuronal culture were carried out as describe previously ***(Zhuang et al. 2019)***. The working concentration for recombinant human Cbln1 (R&D Systems) was 500 ng/ml. GPN (Abcam) was dissolved in DMSO and used at the working concentration of 50 μM. P6-P8 mouse cerebella were cut into small pieces with scissors after the meninges were carefully removed. The tissue was then digested in 5 ml HBSS containing 0.1% Trypsin and 0.04% DNase I in a 37 °C water bath for 15 min before termination addition of 5 ml BME with 10% FBS. Cell suspension was obtained by filtering with sterile cell strainers (40 μm). After centrifuged at 200×g for 5 min, the cell pellets were resuspended in BME supplemented with 5% FBS, 1× GlutaMAX-1, 0.5% glucose and 1× penicillin/streptomycin. The neurons were then plated in PDL-coated cell culture plate and the medium was replaced by maintenance medium supplemented with 1× B27, 1× GlutaMAX-1, 0.5% glucose and 1× penicillin/streptomycin after 4 h.

### Knockdown or overexpression using lentiviral system, RT-qPCR, and Western blotting

The lentiviral knockdown constructs were made using pLKO.1-Puro or pLKO.1-GFP plasmids (Addgene) ***(Zhuang et al. 2019)***. The target sequences of shRNA are as following: *shCbln1*, 5’-GGCTGGAAGTACTCAACCTTC-3’; *shNrxn2*, 5’-CGTTCGTTTATTTCCCTCGAT-3’; *shNrxn2α*, 5’-GGACTTCTGCTGTTCAACTCA-3’; *shNrxn2β*, 5’-CTCCCCCATCACCCGGATTTG-3’; *shCtrl* for pLKO.1-Puro system: 5′-GCATCAAGGTGAACTTCAAGA-3′; *shCtrl* for pLKO.1-GFP system: 5′-GCATAAACCCGCCACTCATCT-3′. RT-qPCR was performed as previously reported***(Zhuang et al. 2019)***. Primers used in qPCR are as following: *mCbln1*, 5’-CCGAGATGAGTAATCGCACCA-3’ and 5’-TCAACATGAGGCTCACCTGGATG-3’; *mNrxn2*, 5’-TACCCGGCAGGAAACTTTGA-3’ and 5’-CCCCCTATCTTGATGGCAGC-3’; *mNrxn2α*, 5’-CTCAAGTCTGGGGCTGTCTG-3’ and 5’-ATAGCGTGTCCAATCCCTGC-3’; *mNrxn2β*, 5’-GATGGATCCAGGCTTCACGG-3’ and 5’-GAAGGAAAACCAGAGCCCGA-3’; *mGapdh*: *5’*-TTGTCAGCAATGCATCCTGCACCACC-3’ and 5’-CTGAGTGGCAGTGATGGCATGGAC-3’.

The coding sequence of mouse *Cbln1* was cloned from E11.5 mouse spinal cord cDNA with primers 5’-CCGGAGGCGCGATGCT-3’ and 5’-ATTCCCGATACGTGCCAG-3’, and lenti viral expression construct was constructed using the pHBLV-CMV-MCS-3×Flag-EF1-Zsgreen1-T2A-Puro backbone (Hanbio). After infection with the lenti virus, the COS7 cell line stably expressing Cbln1 was acquired after multiple rounds of selection using puromycin. Expression of Cbln1-FLAG in cell pellets and supernatant was validated by Western Blotting (WB) following the standard protocols. The dilutions and sources of antibodies used in WB are as following: Cbln1 (1:100, Abclonal), FLAG (1:1000, Beyotime), β-actin (1:10000, Abcam).

### Identification of the differentially expressed genes in the dorsal spinal cord of mouse embryos

*Wnt1-cre* and *Rosa26mTmG* mice were mated to generate *Wnt1-Cre,Rosa26mTmG* embryos. E10.5, E11.5 and E12.5 embryos were collected and dissected, and dorsal spinal cords were dissociated and GFP^+^ neurons were purified using FACS. RNAs were prepared from these purified neurons and the expression profiling was carried out using microarray analysis with GeneChip® Mouse Exon 1.0 ST Array (Affymetrix) following the manufacturer’s manual.

### In situ hybridization

*In situ* hybridization using DIG-labeled RNA probes was carried out on sections from mouse or chick tissue sections following a previously reported protocol ***(Ji and Jaffrey 2012)***. The primers used for PCR in cloning the templates for generating RNA probes are as following (all mouse clones except indicated): *Cbln1*, 5’-CCAAGACGTGACACGCGAGG-3’ and 5’-CAGTAAGTGGCAGGGTTCAG-3’; chick *Cbln1*, 5’-GAGAAGACGCCGCTCAGGTGT-3’ and 5’-CGGGTTGATTTGCGGTCCTTC-3’; *Nrxn1*, 5’-CAGGGAATGCGATCAGGAGG-3’ and 5’-AGACTTCTTCTCTGGCACGC-3’; *Nrxn2*, 5’-TCACAGCCCTGGGTTGATTT-3’ and 5’-AGCAGCGACACACACAAAAG-3’; *Nrxn3*, 5’-GTGAGATGGGGTGTACCACG-3’ and 5’-ACACACACACTGGTCAGAACC-3’; *GluD1*, 5’-CATTGGCCTCCTTCTTGCCT-3’ and 5’-GAGGTGCCATGAGAGGTGTC-3’; *GluD2*, 5’-GCCCCTACCGTGATGTCTTT-3’ and 5’-GTCAATGTCCAGAGGGGTCA-3’; *Nrxn2α*, 5’-GCAGGGATTGGACACGCTAT-3’ and 5’-GAACTGTGACTGCCTACCCC-3’. The template for *Nrxn2β*-specific RNA probe was synthesized (5’-TGAGGGGGGACCCCTAGCCGCCCGCGATGGATCCAGGCTTCACGGACCTTGGCCTTCCCGCTGCGCGTACCCCGGATTCCCCGGCGGGATCCAGTTGATTTGCTTGGCTCCGGACTGAGGCTCGGGCTCTGGTTTTCCTTCGCTTCACCCCTACCCCCCTCTCGGAGCTCGCAACCGGAGGGGGGCTTT-3’) and cloned to pUC57 (Sangon). RNA probes were transcribed *in vitro* using DIG RNA Labeling Kit (SP6/T7) (Roche). Anti-Digoxigenin-AP and NBT/BCIP Stock Solution were also from Roche. *In situ* hybridization images were collected with Axio Imager A2 (Zeiss) or TissueFAXS Cytometer (TissueGnostics).

### Axon guidance assay using co-culture of COS7 cell aggregates with DCN or retinal explants

Aggregates of COS7 cells stably expressing Cbln1 were prepared by resuspending cells in rat tail collagen gel and then placed into 24-well glass bottom plates (Nest). DCN explants were dissected from E11 mouse embryos, immersed in collagen gel and placed 200-400 µm away from the COS7 aggregates. Explants and cell aggregates were co-cultured for 40-48 h in neurobasal medium supplemented with B27, GlutaMAX-1 and penicillin/streptomycin. Similarly, retinal explants were dissected from E14.5 mouse retinas and co-cultured with COS7 aggregates for around 30 h, with the culture medium as following: Neurobasal medium mixed with DMEM/F12 (1:1), supplemented with B27, N21 MAX Media Supplement, GlutaMAX-1 and penicillin/streptomycin.

### DiI tracing of axons

DiI tracing of commissural axons in the spinal cord was performed as previously reported ***(Zhuang et al. 2019)***. DiI labeling of cerebellar parallel fibers was performed as previously described ***(Yamasaki et al. 2001)***. DiI tracing of optic nerves was performed as previously reported ***(Peng et al. 2018)***.

### CTB labelling of optic nerve

To label RGC axon terminals in P4 mouse brain, RGC axons were anterogradely labeled by CTB (Cholera Toxin Subunit B) conjugated with Alexa Fluor™ 555 (Invitrogen, C34776) through intravitreal injection 48 hr before sacrifice. After PFA perfusion, the brains were fixed with 4% PFA in 0.1 M PB overnight, dehydrated with 15% sucrose and 30% sucrose in 0.1 M PB overnight at 4°C sequentially, embedded with O.C.T. for coronal section, and cryosectioned at 12 μm with Leica CM1950 Cryostat. The images were captured on Tissue Genostics with identical settings for each group in the same experiment with the TissueFAXS 7.0 software.

### Immunostaining and immunofluorescence

Immunostaining of tissue sections and immunofluorescence (IF) of cultured DCN explants and neurons were done as previously described ***(Zhuang et al. 2019)***. The spinal cord open-books were prepared similarly as DiI tracing, and their immunostaining was performed with similar procedures as tissues sections except that all incubation and washing were done in 24-well plates and the open-books were mounted onto slides and covered with cover slips before confocal imaging. For IF of axon surface Cbln1 in cultured DCN neurons after GPN treatment, the permeabilization step was omitted and there was no Triton x-100 in antibody incubation buffers. Immunostaining of tissue sections from the cerebellum and the optic chiasm were done as previously described ***(Hirai et al. 2005; Peng et al. 2018)***. The dilutions and sources of antibodies used in immunostaining and immunofluorescence are as following: Cbln1 (1:1000, Abcam), Cbln1 (1:50, Frontier Institute), Lhx2 (1:500, Abcam), Alcam (1:200, R&D Systems), Lhx9 (1:50, Santa Cruz Biotechnology), Robo3 (1:500, R&D Systems), Tag1 (1:200, R&D Systems), NFM (1:1000, Cell Signaling Technology), GFP (1:1000, Abcam), Isl1/2 (1:500, DSHB), Nrxn2 (1:200, Abcam), Calbindin (1:200, Swant), NeuN (1:500, Cell Signaling Technology), Brn3a (1:200, Millipore). Alexa Fluor-conjugated secondary antibodies (Thermo) were used at 1:1000 (555) or 1:500 (488). Fluorescent images were acquired using laser-scanning confocal microscopes Nikon A1R with NIS software, Leica SP8 with LASX software, or Zeiss LSM 800 with Zen software. All images were collected with identical settings for each group in the same experiment. Quantification of immunofluorescence signals was performed using ImageJ.

### Statistical analysis

Statistical analysis was performed with GraphPad Prism 7.0. Most of our data are represented as the box and whisker plots unless otherwise specified in the figure legends, and the settings are: 25–75 percentiles (boxes), minimum and maximum (whiskers), and medians (horizontal lines). Unpaired Student’s t test was performed for comparison between two groups and ANOVA with Tukey’s multiple comparison test was performed to the comparison of three or more groups. * indicates statistically significant: \**p* < 0.05, \*\**p* < 0.01, \*\*\**p* < 0.001, \*\*\*\**p* <0.0001.

## Acknowledgements

We thank Esther T. Stoeckli for the pMath1-eGFP-miRNA vector, members of Ji and Jaffrey laboratories for help, technical support, and comments on the manuscript. This work was supported by National Natural Science Foundation of China (31871038 and 32170955 to S.-J.J.), Shenzhen-Hong Kong Institute of Brain Science-Shenzhen Fundamental Research Institutions (2021SHIBS0002, 2019SHIBS0002), High-Level University Construction Fund for Department of Biology (internal grant no. G02226301), Science and Technology Innovation Commission of Shenzhen Municipal Government (ZDSYS20200811144002008), and NIH (R35NS111631 to S.R.J.).

## Author contributions

S.-J.J. and S.R.J. conceived the project and designed experiments; S.-J.J. performed screening and identification of DEGs in DCNs; P.H. and Y.S. performed and analyzed most of the experiments with help of Z.Y., M.Z. and Q.W.; X.L., C.Y., and J.Z. performed experiments using chick embryos; S.-J.J., S.R.J., and P.H. wrote the manuscript with editing and input from other authors.

## Ethics

All experiments using mice were carried out following the animal protocols approved by the Laboratory Animal Welfare and Ethics Committee of Southern University of Science and Technology (approval numbers: SUSTC-JY2017004, SUSTech-JY202102081).

## Data availability statement

The microarray data has been deposited to the Gene Expression Omnibus (GEO) with accession number GSE169448.

---

**Figure 1—figure supplement 1.**
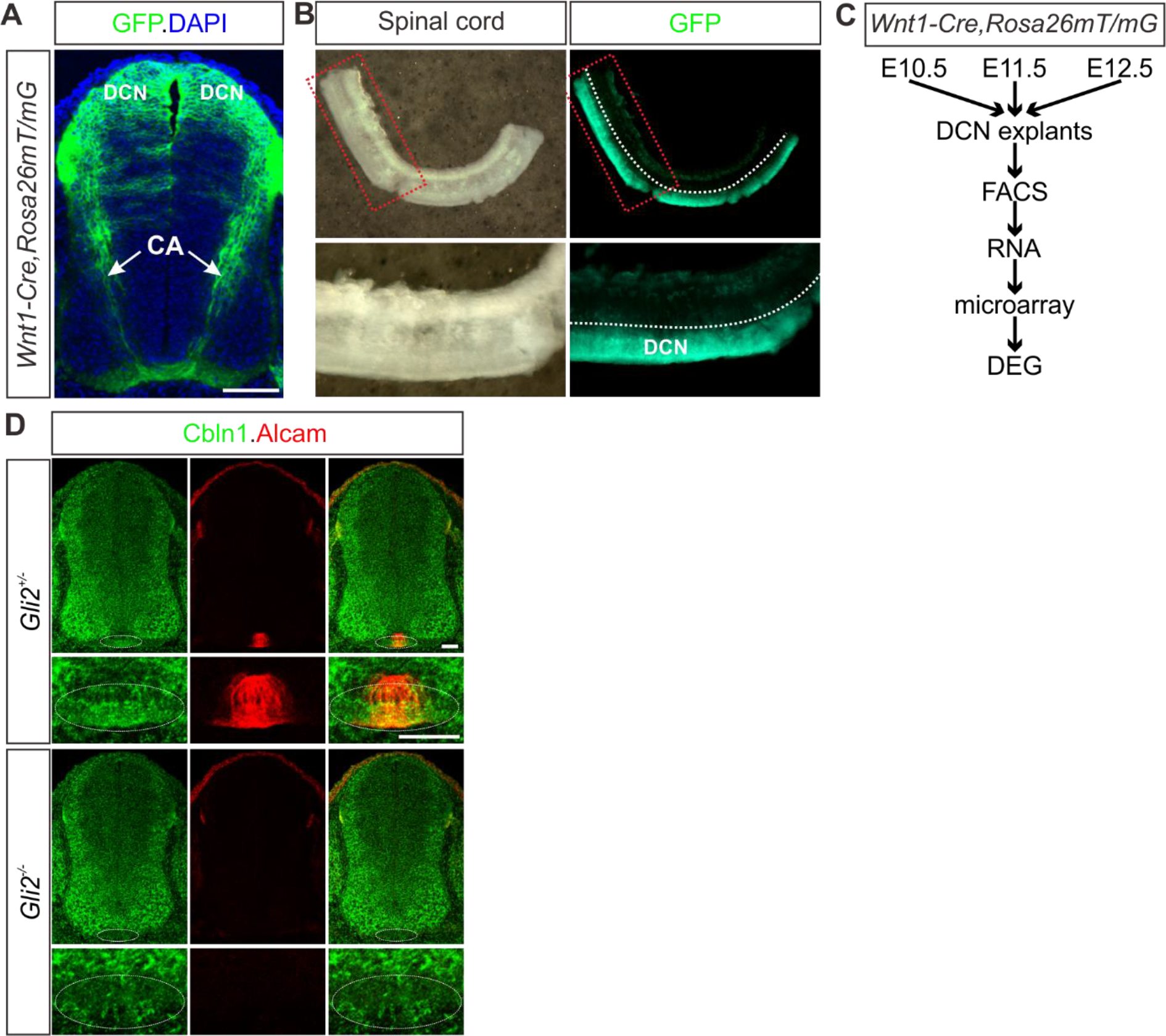
Screening of the differentially expressed genes in the mouse embryonic dorsal spinal cord. **(A)** The embryonic dorsal spinal neurons were genetically labeled with eGFP by crossing *Wnt1-cre* with *Rosa26mTmG* mice. Immunofluorescence of cross-sections of E11.5 spinal cord was shown. DCN, dorsal commissural neurons. CA, commissural axons. Scale bar, 100 μm. **(B)** The dissected E11.5 spinal cords were shown in both bright-field and fluorescent images. The regions in the red dotted boxes were shown with higher magnification in the lower images. The dotted white line indicates where to cut and separate dorsal and ventral spinal cord. **(C)** The scheme showing the procedures for identifying the differentially expressed genes in the mouse embryonic dorsal spinal cord. **(D)** Co-immunostaining of Cbln1 with Alcam in spinal cord cross-sections of *Gli2* KO and its littermate control embryos at E11.5. The circled areas indicate the expression and loss of Cbln1 in control and *Gli2* KO embryos, respectively. Scale bar, 50 μm.

**Figure 2—figure supplement 1.**
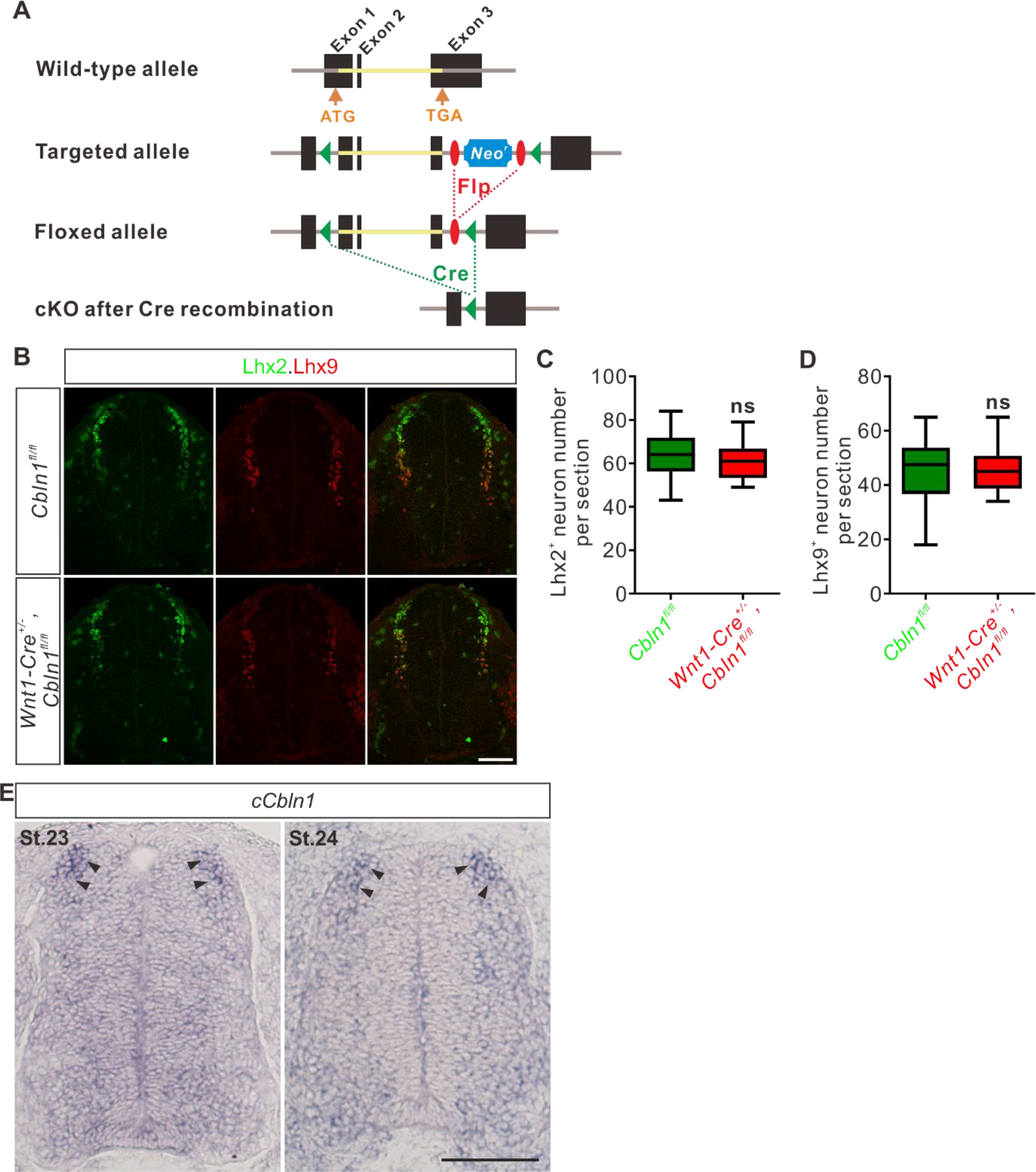
DCN-specific cKO of *Cbln1* does not affect neurogenesis and patterning of spinal DCNs. **(A)** Schematic drawings showing the generation of *Cbln1* cKO. The coding sequence of *Cbln1* is deleted after Cre-mediated recombination. **(B)** Lhx2 and Lhx9 immunostaining in E11.5 spinal cord indicated that *Cbln1* cKO in DCNs does not disturb neurogenesis and patterning of spinal DCNs. Scale bar, 100 μm. **(C, D)** Quantification of Lhx2^+^ and Lhx9^+^ neurons in (**B**) showed that their neurogenesis or patterning is not affected in *Cbln1* cKO. All data are represented as box and whisker plots: *Cbln1^fl/fl^* (*n* = 20 sections) vs *Wnt1-Cre^+/-^,Cbln1^fl/fl^* (*n* = 16 sections); ns, not significant; *p* = 0.46 for Lhx2^+^ neurons in (**C**); *p* = 0.99 for Lhx9^+^ neurons in (**D**); by unpaired Student’s *t* test. **(E)** *In situ* hybridization of *cCbln1* in spinal cord cross-sections of St.23-24 chick embryos. Arrowheads indicate the expression of *cCbln1* in DCNs. Scale bar, 50 μm.

**Figure 3—figure supplement 1.**
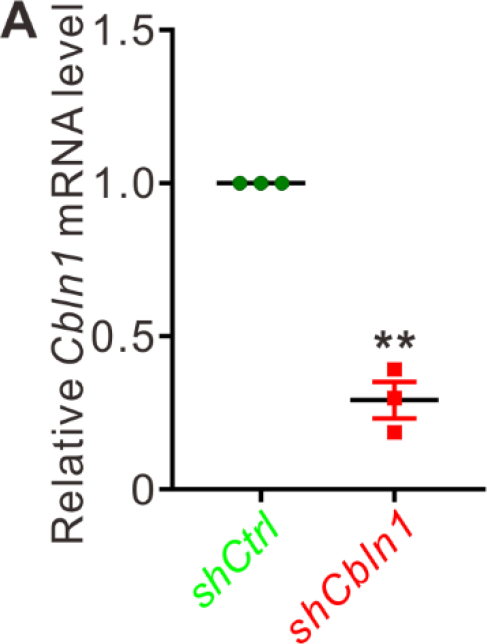
Knockdown of mCbln1 using lentiviral shRNA. **(A)** Validation of knockdown of *Cbln1* using lentiviral *shCbln1*. Since Cbln1 is abundant in cerebellar granule cells, we prepared dissociated cerebellar granule cells from P8 mouse pups and cultured *in vitro* to test the knockdown efficiency of *shCbln1*. *Cbln1* mRNA levels were measured by RT-qPCR after lentiviral shRNA infection. Data are mean ± SEM. and represented as dot plots: \**\*p* = 0.0070; by unpaired Student’s *t* test.

**Figure 4—figure supplement 1.**
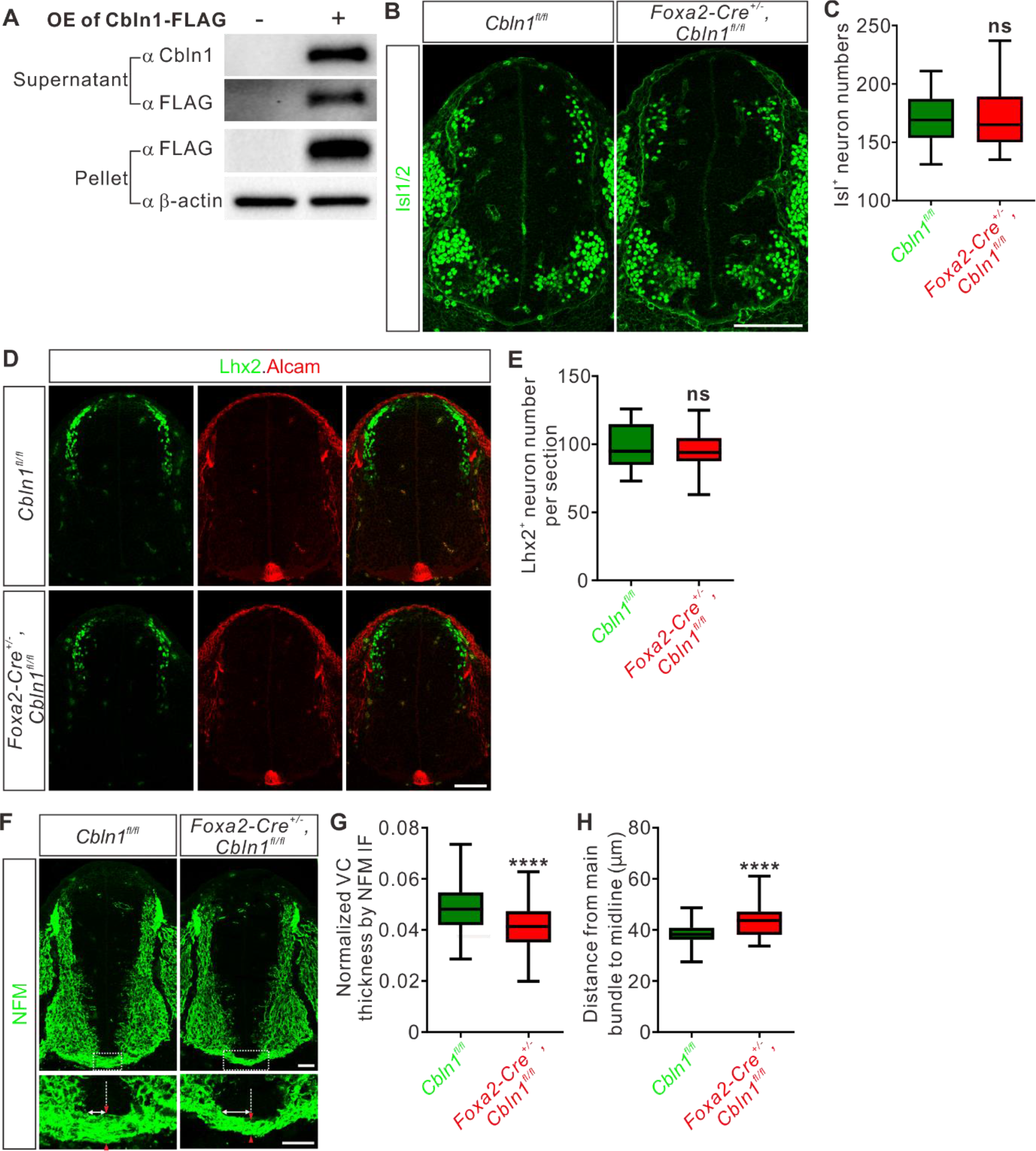
Floor plate-specific *Cbln1* cKO does not disturb neural patterning or neurogenesis in the developing spinal cord. **(A)** Overexpression and secretion of Cbln1 tagged by FLAG in COS7 cells were validated by WB. **(B, C)** Isl1/2 immunostaining showed normal patterning of spinal cord in the floor plate-specific *Cbln1* cKO embryos (**B**). Isl1/2 marks different interneurons and motor neurons in spinal cord. Scale bar, 100 μm. The data for quantification of Isl1/2^+^ neuron numbers are represented as box and whisker plots (**C**): *Cbln1^fl/fl^* (*n* = 42 sections) vs *Foxa2-Cre^+/-^,Cbln1^fl/fl^* (*n* = 49 sections); ns, not significant (*p* = 0.93); by unpaired Student’s *t* test. **(D, E)** The floor plate-specific cKO of Cbln1 does not disturb DCN neurogenesis, spinal cord patterning, or floor plate development. Lhx2 and Alcam immunostainings of E11.5 spinal cord were used to mark dI1 commissural neurons and floor plate, respectively (**D**). Scale bar, 100 μm. The data for quantification of Lhx2^+^ neuron numbers are represented as box and whisker plots (**E**): *Cbln1^fl/fl^* (*n* = 16 sections) vs *Foxa2-Cre^+/-^,Cbln1^fl/fl^* (*n* = 35 sections); ns, not significant (*p* = 0.59); by unpaired Student’s *t* test. **(F)** The axon guidance defects of pre-crossing commissural axons were observed by NFM immunostaining in floor plate-specific *Cbln1* cKO and control embryos at E11.5. Higher magnification views of the FP region in the white dotted boxes are also shown (bottom). The pair of red arrowheads denotes the thickness of the ventral commissure (VC). The double-arrowed line measures the distance between the point of intersection (of the main pre-crossing commissural axon bundle with the ventral edge of spinal cord) and the midline (indicated by the dotted line). Scale bars, 50 μm. **(G, H)** Quantification of the VC thickness and the distance from the main bundle intersection point to the midline. All data are represented as box and whisker plots: *Cbln1^fl/fl^* (*n* = 49 sections) vs *Foxa2-Cre^+/-^,Cbln1^fl/fl^* (*n* = 74 sections), \*\*\**\*p* = 3.86E-05 for (**G**), \*\*\**\*p* = 1.35E-07 for (**H**), by unpaired Student’s *t* test.

**Figure 5—figure supplement 1.**
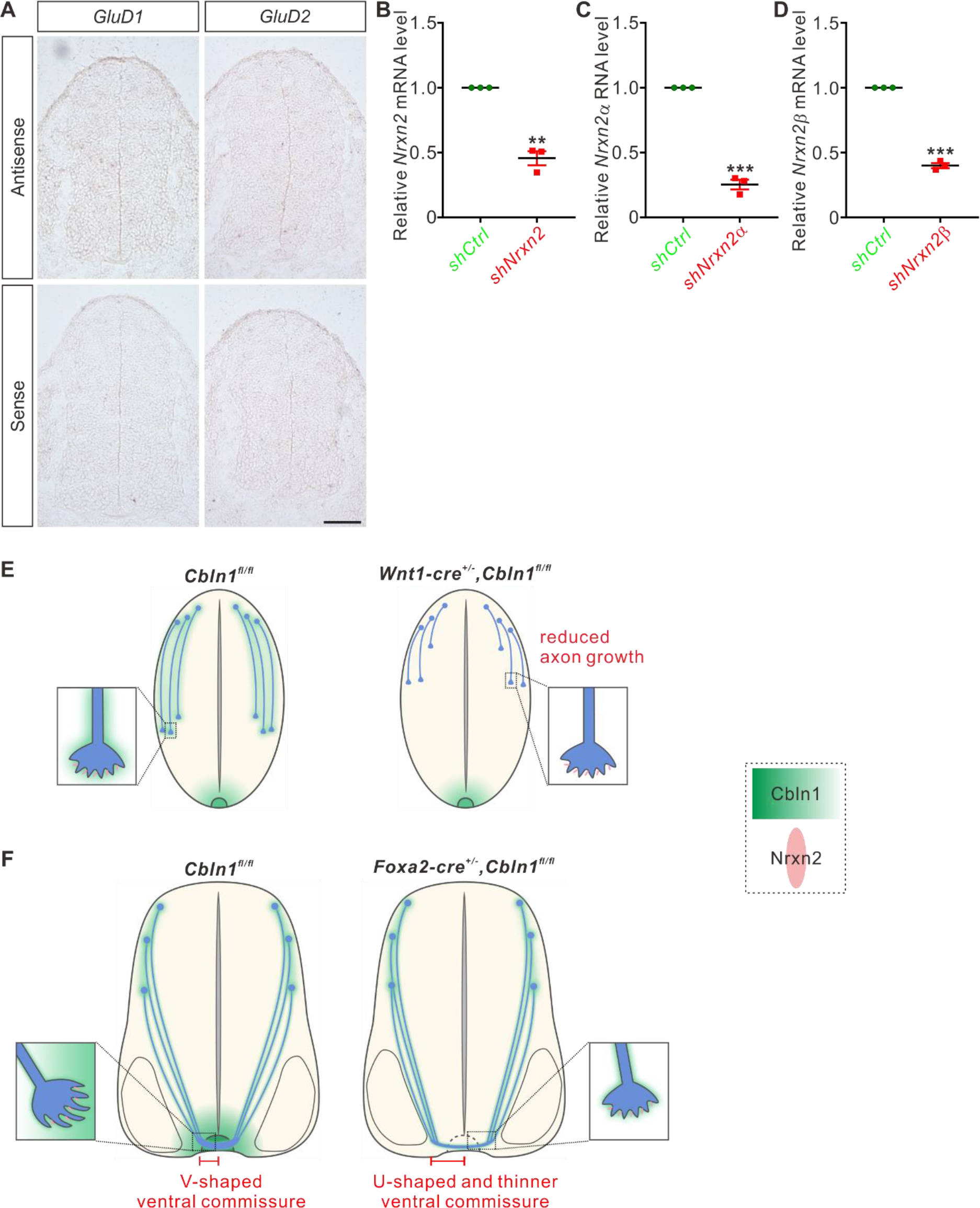
Working models for cell-autonomous and non-cell-autonomous Cbln1 in the developing spinal cord. **(A)** *GluD1 or GLuD2* mRNA was not detected in E11.5 spinal cord cross-sections by *in situ* hybridization. Scale bar, 100 μm. **(B-D)** Validation of knockdown by shRNAs. Dissociated cerebellar granule cells from P8 mouse pups were cultured and lentiviral shRNAs were infected. Significant knockdown was achieved by shRNAs against *Nrxn2*, *Nrxn2α* and *Nrxn2β*, respectively. RT-qPCR data are mean ± SEM. and represented as dot plots: \**\*p* = 0.0084 for (**B**); \*\**\*p* = 0.00010 for (**C**); \*\**\*p* = 0.00020 for (**D**); by unpaired Student’s *t* test. **(E)** Working model for the stimulation of commissural axon growth by the cell-autonomous Cbln1. In the pre-crossing commissural axons, Cbln1 is expressed cell-autonomously by the dorsal commissural neurons and axons. Commissural axon growth cone-secreted Cbln1 works back to itself in an autocrine manner and binds to Nrxn2 receptors to stimulate commissural axon growth. In the DCN-specific *Cbln1* cKO embryos, commissural axon growth is reduced compared with their littermate controls. **(F)** Working model for the attraction of commissural axon growth toward midline by the non-cell-autonomous, floor plate-derived Cbln1. When commissural axons approach the midline, the floor plate-derived Cbln1 attracts commissural axons to the midline which is also mediate by Nrxn2 receptors. In the floor plate-specific *Cbln1* cKO embryos, commissural axon guidance in the midline crossing is impaired, resulting in a U-shaped and thinner ventral commissure compared with the V-shaped and thick ventral commissures in the littermate control embryos.

**Figure 6—figure supplement 1.**
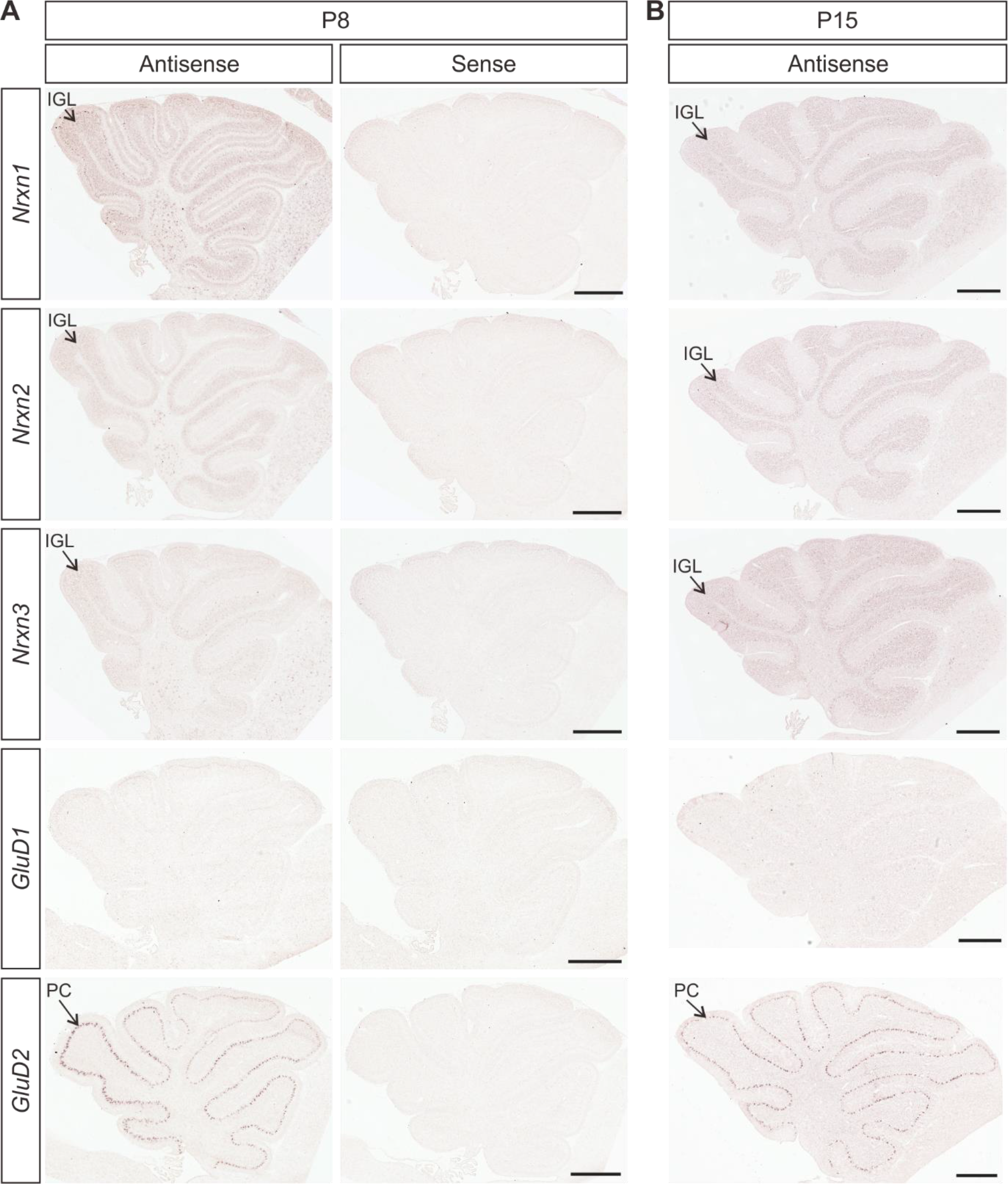
Expression of Cbln1 receptors in the developing cerebellum. **(A, B)** *In situ* hybridization of *Nrxn1, Nrxn2, Nrxn3, GluD1* and *GluD2* in cerebella at P8 (**A**) and P15 (**B**). *Nrxn1, Nrxn2* and *Nrxn3* mRNAs were detected in the inner granule layer (IGL). *GluD2* mRNA was highly and specifically expressed in the Purkinje cells (PCs) while *GluD1* mRNA was not detected in the cerebellum at these stages. Scale bars, 500 μm.

**Figure 7—figure supplement 1.**
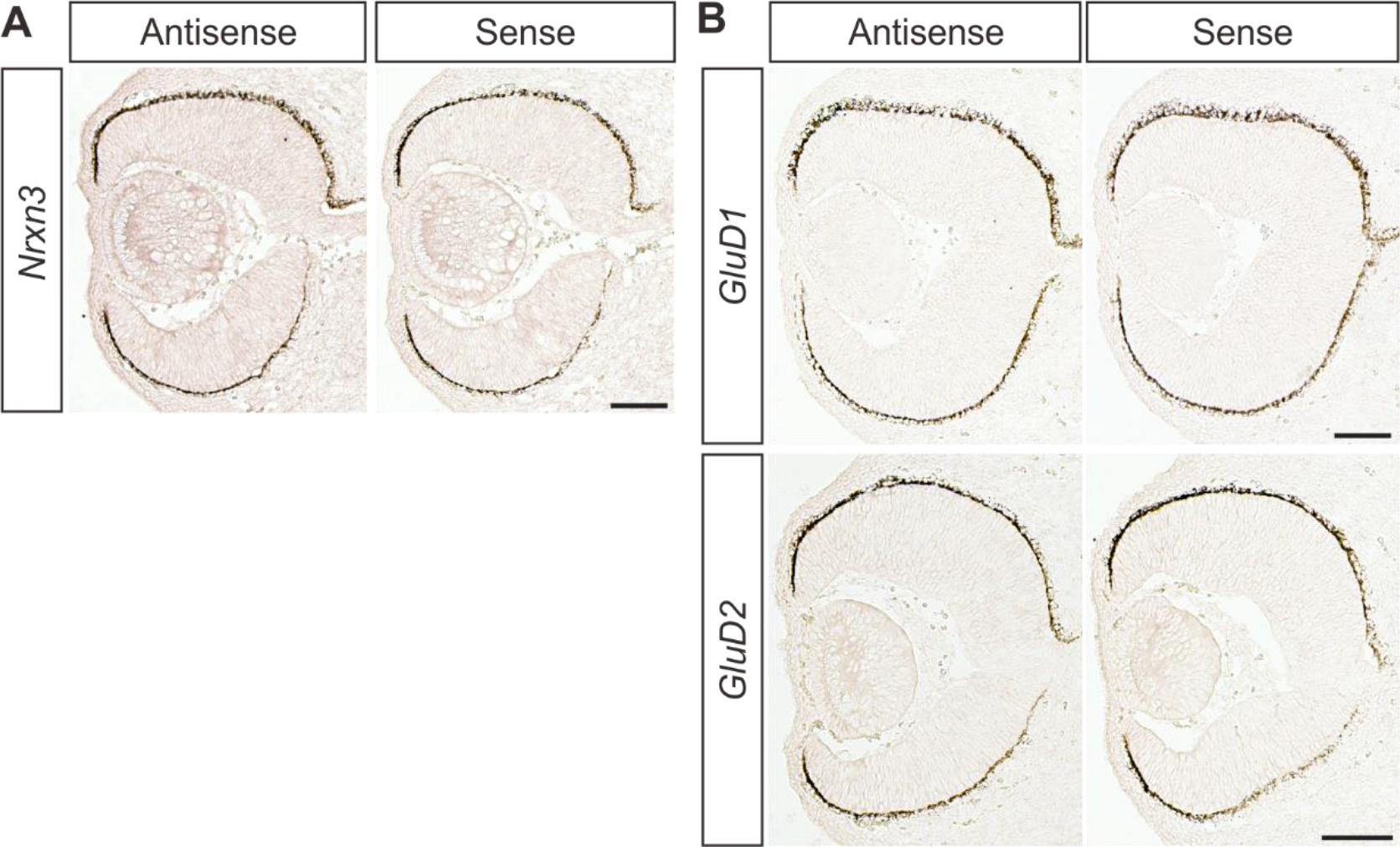
*In situ* hybridization of Cbln1 receptors in the developing retina. **(A, B)** *In situ* hybridization of *Nrxn3* (**A**), and *GluD1* and *GluD2* (**B**) in E13 retina. *Nrxn3*, *GluD1* or *GluD2* mRNA was not detected in the developing retina. Scale bars, 100 μm.

**Supplementary file 1. Differentially expressed genes in the dorsal spinal cord of mouse embryos.**

See the separate Excel data set

## References

1. Augsburger A, Schuchardt A, Hoskins S, Dodd J, Butler S. 1999. BMPs as mediators of roof plate repulsion of commissural neurons. Neuron 24:127–141. PMID: 10677032

2. Bai CB, Joyner AL. 2001. Gli1 can rescue the in vivo function of Gli2. Development 128:5161–5172. PMID: 11748151

3. Biederer T, Sudhof TC. 2001. CASK and protein 4.1 support F-actin nucleation on neurexins. J Biol Chem 276:47869–47876. DOI: https://doi.org/10.1074/jbc.M105287200, PMID: 11604393

4. Biederer T, Südhof TC. 2000. Mints as adaptors. Direct binding to neurexins and recruitment of munc18. J Biol Chem 275:39803-39806. DOI: https://doi.org/10.1074/jbc.C000656200, PMID: 11036064

5. Brose K, Bland KS, Wang KH, Arnott D, Henzel W, Goodman CS, Tessier-Lavigne M, Kidd T. 1999. Slit proteins bind Robo receptors and have an evolutionarily conserved role in repulsive axon guidance. Cell 96:795–806. PMID: 10102268

6. Butler SJ, Dodd J. 2003. A role for BMP heterodimers in roof plate-mediated repulsion of commissural axons. Neuron 38:389–401. PMID: 12741987

7. Cerrato V, Mercurio S, Leto K, Fucà E, Hoxha E, Bottes S, Pagin M, Milanese M, Ngan CY, Concina G, Ottolenghi S, Wei CL, Bonanno G, Pavesi G, Tempia F, Buffo A, Nicolis SK. 2018. Sox2 conditional mutation in mouse causes ataxic symptoms, cerebellar vermis hypoplasia, and postnatal defects of Bergmann glia. Glia 66:1929–1946. DOI: https://doi.org/10.1002/glia.23448, PMID: 29732603

8. Charron F, Stein E, Jeong J, McMahon AP, Tessier-Lavigne M. 2003. The morphogen sonic hedgehog is an axonal chemoattractant that collaborates with netrin-1 in midline axon guidance. Cell 113:11–23. PMID: 12679031

9. Chen Z, Gore BB, Long H, Ma L, Tessier-Lavigne M. 2008. Alternative splicing of the Robo3 axon guidance receptor governs the midline switch from attraction to repulsion. Neuron 58:325–332. DOI: https://doi.org/10.1016/j.neuron.2008.02.016, PMID: 18466743

10. Colak D, Ji SJ, Porse BT, Jaffrey SR. 2013. Regulation of axon guidance by compartmentalized nonsense-mediated mRNA decay. Cell 153:1252–1265. DOI: https://doi.org/10.1016/j.cell.2013.04.056, PMID: 23746841

11. Danielian PS, Muccino D, Rowitch DH, Michael SK, McMahon AP. 1998. Modification of gene activity in mouse embryos in utero by a tamoxifen-inducible form of Cre recombinase. Curr Biol 8:1323–1326. PMID: 9843687

12. Fossati M, Assendorp N, Gemin O, Colasse S, Dingli F, Arras G, Loew D, Charrier C. 2019. Trans-Synaptic Signaling through the Glutamate Receptor Delta-1 Mediates Inhibitory Synapse Formation in Cortical Pyramidal Neurons. Neuron 104:1081–1094 e1087. DOI: https://doi.org/10.1016/j.neuron.2019.09.027, PMID: 31704028

13. Harada H, Farhani N, Wang XF, Sugita S, Charish J, Attisano L, Moran M, Cloutier JF, Reber M, Bremner R, Monnier PP. 2019. Extracellular phosphorylation drives the formation of neuronal circuitry. Nat Chem Biol 15:1035–1042. DOI: https://doi.org/10.1038/s41589-019-0345-z, PMID: 31451763

14. Hata Y, Butz S, Südhof TC. 1996. CASK: a novel dlg/PSD95 homolog with an N-terminal calmodulin-dependent protein kinase domain identified by interaction with neurexins. J Neurosci 16:2488–2494. DOI: https://doi.org/10.1523/jneurosci.16-08-02488.1996, PMID: 8786425

15. Hernandez-Enriquez B, Wu Z, Martinez E, Olsen O, Kaprielian Z, Maness PF, Yoshida Y, Tessier-Lavigne M, Tran TS. 2015. Floor plate-derived neuropilin-2 functions as a secreted semaphorin sink to facilitate commissural axon midline crossing. Genes Dev 29:2617–2632. DOI: https://doi.org/10.1101/gad.268086.115, PMID: 26680304

16. Hirai H, Pang Z, Bao D, Miyazaki T, Li L, Miura E, Parris J, Rong Y, Watanabe M, Yuzaki M, Morgan JI. 2005. Cbln1 is essential for synaptic integrity and plasticity in the cerebellum. Nat Neurosci 8:1534–1541. DOI: https://doi.org/10.1038/nn1576, PMID: 16234806

17. Ibata K, Kono M, Narumi S, Motohashi J, Kakegawa W, Kohda K, Yuzaki M. 2019. Activity-Dependent Secretion of Synaptic Organizer Cbln1 from Lysosomes in Granule Cell Axons. Neuron 102:1184–1198.e1110. DOI: https://doi.org/10.1016/j.neuron.2019.03.044, PMID: 31072786

18. Islam SM, Shinmyo Y, Okafuji T, Su Y, Naser IB, Ahmed G, Zhang S, Chen S, Ohta K, Kiyonari H, Abe T, Tanaka S, Nishinakamura R, Terashima T, Kitamura T, Tanaka H. 2009. Draxin, a repulsive guidance protein for spinal cord and forebrain commissures. Science 323:388–393. DOI: https://doi.org/10.1126/science.1165187, PMID: 19150847

19. Ito-Ishida A, Miyazaki T, Miura E, Matsuda K, Watanabe M, Yuzaki M, Okabe S. 2012. Presynaptically released Cbln1 induces dynamic axonal structural changes by interacting with GluD2 during cerebellar synapse formation. Neuron 76:549–564. DOI https://doi.org/10.1016/j.neuron.2012.07.027, PMID: 23141067

20. Ji SJ, Jaffrey SR. 2012. Intra-axonal translation of SMAD1/5/8 mediates retrograde regulation of trigeminal ganglia subtype specification. Neuron 74:95–107. DOI: https://doi.org/10.1016/j.neuron.2012.02.022, PMID: 22500633

21. Ji SJ, Periz G, Sockanathan S. 2009. Nolz1 is induced by retinoid signals and controls motoneuron subtype identity through distinct repressor activities. Development 136:231–240. DOI: https://doi.org/10.1242/dev.028043, PMID: 19056829

22. Kidd T, Bland KS, Goodman CS. 1999. Slit is the midline repellent for the robo receptor in Drosophila. Cell 96:785–794. PMID: 10102267

23. Lyuksyutova AI, Lu CC, Milanesio N, King LA, Guo N, Wang Y, Nathans J, Tessier-Lavigne M, Zou Y. 2003. Anterior-posterior guidance of commissural axons by Wnt-frizzled signaling. Science 302:1984–1988. DOI: https://doi.org/10.1126/science.1089610, PMID: 14671310

24. Mains RE, Kiraly DD, Eipper-Mains JE, Ma XM, Eipper BA. 2011. Kalrn promoter usage and isoform expression respond to chronic cocaine exposure. BMC neuroscience 12:20. DOI: https://doi.org/10.1186/1471-2202-12-20, PMID: 21329509

25. Mason C, Slavi N. 2020. Retinal Ganglion Cell Axon Wiring Establishing the Binocular Circuit. Annu Rev Vis Sci. DOI: https://doi.org/10.1146/annurev-vision-091517-034306, PMID: 32396770

26. Matsuda K, Miura E, Miyazaki T, Kakegawa W, Emi K, Narumi S, Fukazawa Y, Ito-Ishida A, Kondo T, Shigemoto R, Watanabe M, Yuzaki M. 2010. Cbln1 is a ligand for an orphan glutamate receptor delta2, a bidirectional synapse organizer. Science 328:363–368. DOI: https://doi.org/10.1126/science.1185152, PMID: 20395510

27. McCormick LE, Gupton SL. 2020. Mechanistic advances in axon pathfinding. Curr Opin Cell Biol 63:11–19. DOI: https://doi.org/10.1016/j.ceb.2019.12.003, PMID: 31927278

28. McCurdy EP, Chung KM, Benitez-Agosto CR, Hengst U. 2019. Promotion of Axon Growth by the Secreted End of a Transcription Factor. Cell Rep 29:363–377.e365. DOI: https://doi.org/10.1016/j.celrep.2019.08.101, PMID: 31597097

29. Moreno-Bravo JA, Roig Puiggros S, Mehlen P, Chedotal A. 2019. Synergistic Activity of Floor-Plate- and Ventricular-Zone-Derived Netrin-1 in Spinal Cord Commissural Axon Guidance. Neuron 101:625–634.e623. DOI: https://doi.org/10.1016/j.neuron.2018.12.024, PMID: 30661739

30. Muguruma K, Nishiyama A, Ono Y, Miyawaki H, Mizuhara E, Hori S, Kakizuka A, Obata K, Yanagawa Y, Hirano T, Sasai Y. 2010. Ontogeny-recapitulating generation and tissue integration of ES cell-derived Purkinje cells. Nat Neurosci 13:1171–1180. DOI: https://doi.org/10.1038/nn.2638, PMID: 20835252

31. Mukherjee K, Sharma M, Urlaub H, Bourenkov GP, Jahn R, Südhof TC, Wahl MC. 2008. CASK Functions as a Mg2+-independent neurexin kinase. Cell 133:328–339. DOI: https://doi.org/10.1016/j.cell.2008.02.036, PMID: 18423203

32. Muzumdar MD, Tasic B, Miyamichi K, Li L, Luo L. 2007. A global double-fluorescent Cre reporter mouse. Genesis 45:593–605. DOI: https://doi.org/10.1002/dvg.20335, PMID: 17868096

33. Nawabi H, Briancon-Marjollet A, Clark C, Sanyas I, Takamatsu H, Okuno T, Kumanogoh A, Bozon M, Takeshima K, Yoshida Y, Moret F, Abouzid K, Castellani V. 2010. A midline switch of receptor processing regulates commissural axon guidance in vertebrates. Genes Dev 24:396–410. DOI: https://doi.org/10.1101/gad.542510, PMID: 20159958

34. Okada A, Charron F, Morin S, Shin DS, Wong K, Fabre PJ, Tessier-Lavigne M, McConnell SK. 2006. Boc is a receptor for sonic hedgehog in the guidance of commissural axons. Nature 444:369–373. DOI: https://doi.org/10.1038/nature05246, PMID: 17086203

35. Padamsey Z, McGuinness L, Bardo SJ, Reinhart M, Tong R, Hedegaard A, Hart ML, Emptage NJ. 2017. Activity-Dependent Exocytosis of Lysosomes Regulates the Structural Plasticity of Dendritic Spines. Neuron 93:132–146. DOI: https://doi.org/10.1016/j.neuron.2016.11.013, PMID: 27989455

36. Paixao S, Balijepalli A, Serradj N, Niu J, Luo W, Martin JH, Klein R. 2013. EphrinB3/EphA4-mediated guidance of ascending and descending spinal tracts. Neuron 80:1407–1420. DOI: https://doi.org/10.1016/j.neuron.2013.10.006, PMID: 24360544

37. Park EJ, Sun X, Nichol P, Saijoh Y, Martin JF, Moon AM. 2008. System for tamoxifen-inducible expression of cre-recombinase from the Foxa2 locus in mice. Dev Dyn 237:447–453. DOI: https://doi.org/10.1002/dvdy.21415, PMID: 18161057

38. Peng J, Fabre PJ, Dolique T, Swikert SM, Kermasson L, Shimogori T, Charron F. 2018. Sonic Hedgehog Is a Remotely Produced Cue that Controls Axon Guidance Trans-axonally at a Midline Choice Point. Neuron 97:326–340 e324. DOI: https://doi.org/10.1016/j.neuron.2017.12.028, PMID: 29346753

39. Reissner C, Runkel F, Missler M. 2013. Neurexins. Genome Biol 14:213. DOI: https://doi.org/10.1186/gb-2013-14-9-213, PMID: 24083347

40. Sabatier C, Plump AS, Le M, Brose K, Tamada A, Murakami F, Lee EY, Tessier-Lavigne M. 2004. The divergent Robo family protein rig-1/Robo3 is a negative regulator of slit responsiveness required for midline crossing by commissural axons. Cell 117:157–169. PMID: 15084255

41. Stoeckli ET. 2018. Understanding axon guidance: are we nearly there yet? Development 145. DOI: https://doi.org/10.1242/dev.151415, PMID: 29759980

42. Stoeckli ET, Kuhn TB, Duc CO, Ruegg MA, Sonderegger P. 1991. The axonally secreted protein axonin-1 is a potent substratum for neurite growth. The Journal of cell biology 112:449–455. DOI: https://doi.org/10.1083/jcb.112.3.449, PMID: 1991792

43. Sudhof TC. 2017. Synaptic Neurexin Complexes: A Molecular Code for the Logic of Neural Circuits. Cell 171:745–769. DOI: https://doi.org/10.1016/j.cell.2017.10.024, PMID: 29100073

44. Suzuki K, Elegheert J, Song I, Sasakura H, Senkov O, Matsuda K, Kakegawa W, Clayton AJ, Chang VT, Ferrer-Ferrer M, Miura E, Kaushik R, Ikeno M, Morioka Y, Takeuchi Y, Shimada T, Otsuka S, Stoyanov S, Watanabe M, Takeuchi K, Dityatev A, Aricescu AR, Yuzaki M. 2020. A synthetic synaptic organizer protein restores glutamatergic neuronal circuits. Science 369. DOI: https://doi.org/10.1126/science.abb4853, PMID: 32855309

45. Takeo YH, Shuster SA, Jiang L, Hu MC, Luginbuhl DJ, Rülicke T, Contreras X, Hippenmeyer S, Wagner MJ, Ganguli S, Luo L. 2020. GluD2- and Cbln1-mediated competitive interactions shape the dendritic arbors of cerebellar Purkinje cells. Neuron. DOI: https://doi.org/10.1016/j.neuron.2020.11.028, PMID: 33352118

46. Tronche F, Kellendonk C, Kretz O, Gass P, Anlag K, Orban PC, Bock R, Klein R, Schutz G. 1999. Disruption of the glucocorticoid receptor gene in the nervous system results in reduced anxiety. Nat Genet 23:99–103. DOI: https://doi.org/10.1038/12703, PMID: 10471508

47. Uemura T, Lee SJ, Yasumura M, Takeuchi T, Yoshida T, Ra M, Taguchi R, Sakimura K, Mishina M. 2010. Trans-synaptic interaction of GluRdelta2 and Neurexin through Cbln1 mediates synapse formation in the cerebellum. Cell 141:1068–1079. DOI: https://doi.org/10.1016/j.cell.2010.04.035, PMID: 20537373

48. Wilson NH, Stoeckli ET. 2013. Sonic hedgehog regulates its own receptor on postcrossing commissural axons in a glypican1-dependent manner. Neuron 79:478–491. DOI: https://doi.org/10.1016/j.neuron.2013.05.025, PMID: 23931997

49. Wilson SI, Shafer B, Lee KJ, Dodd J. 2008. A molecular program for contralateral trajectory: Rig-1 control by LIM homeodomain transcription factors. Neuron 59:413–424. DOI: https://doi.org/10.1016/j.neuron.2008.07.020, PMID: 18701067

50. Wu Z, Makihara S, Yam PT, Teo S, Renier N, Balekoglu N, Moreno-Bravo JA, Olsen O, Chedotal A, Charron F, Tessier-Lavigne M. 2019. Long-Range Guidance of Spinal Commissural Axons by Netrin1 and Sonic Hedgehog from Midline Floor Plate Cells. Neuron 101:635–647.e634. DOI: https://doi.org/10.1016/j.neuron.2018.12.025, PMID: 30661738

51. Yamasaki T, Kawaji K, Ono K, Bito H, Hirano T, Osumi N, Kengaku M. 2001. Pax6 regulates granule cell polarization during parallel fiber formation in the developing cerebellum. Development 128:3133–3144. PMID: 11688562

52. Yasumura M, Yoshida T, Lee SJ, Uemura T, Joo JY, Mishina M. 2012. Glutamate receptor δ1 induces preferentially inhibitory presynaptic differentiation of cortical neurons by interacting with neurexins through cerebellin precursor protein subtypes. Journal of Neurochemistry 121:705–716. DOI: https://doi.org/10.1111/j.1471-4159.2011.07631.x, PMID: 22191730

53. Yu J, She Y, Yang L, Zhuang M, Han P, Liu J, Lin X, Wang N, Chen M, Jiang C, Zhang Y, Yuan Y, Ji SJ. 2021. The m6A Readers YTHDF1 and YTHDF2 Synergistically Control Cerebellar Parallel Fiber Growth by Regulating Local Translation of the Key Wnt5a Signaling Components in Axons. Adv Sci. DOI: https://doi.org/10.1002/advs.202101329,

54. Yuzaki M. 2018. Two Classes of Secreted Synaptic Organizers in the Central Nervous System. Annu Rev Physiol 80:243–262. DOI: https://doi.org/10.1146/annurev-physiol-021317-121322, PMID: 29166241

55. Zhuang M, Li X, Zhu J, Zhang J, Niu F, Liang F, Chen M, Li D, Han P, Ji SJ. 2019. The m6A reader YTHDF1 regulates axon guidance through translational control of Robo3.1 expression. Nucleic acids research 47:4765–4777. DOI: https://doi.org/10.1093/nar/gkz157, PMID: 30843071

56. Zou Y, Stoeckli E, Chen H, Tessier-Lavigne M. 2000. Squeezing axons out of the gray matter: a role for slit and semaphorin proteins from midline and ventral spinal cord. Cell 102:363–375. PMID: 10975526

